# Maladaptive inflammatory signaling in old mice impairs colonic regeneration by promoting a sustained fetal-like epithelial state

**DOI:** 10.1101/2025.07.16.665215

**Authors:** Antonion Korcari, Helen Tauc, Jeff Duggan, Fabien Wehbe, Spyros Darmanis, Zora Modrusan, Jina Yun, Rajita Pappu, David Garfield, Heinrich Jasper, David Castillo-Azofeifa

## Abstract

Aging is associated with a decline in the regenerative capacity of many tissues. Central to this decline is a complex interplay between inflammation and stem cell function. How these two processes are linked and influence regenerative capacity remains unclear. Here, we undertake a comprehensive assessment of age-related changes in the mouse colon at single-cell resolution. A survey of immune and epithelial compartments revealed a hyperactivated inflammatory state in the colon of old mice characterized by the induction of an interferon γ (IFNγ) response signature in immune cells. This does not result in increased inflammation under homeostasis, but triggers a disproportionate inflammatory response, disrupting regeneration after challenge with the enteropathogen *Citrobacter rodentium*. Colons of old mice exhibit higher production of IFNγ by T and innate lymphoid cells (ILCs) that are associated with reduced Lgr5^+^ stem cells and decreased epithelial proliferation. Interestingly, we find aged intestinal epithelial cells to be hypersensitive to IFNγ signaling, inducing a regeneration-associated fetal-like gene expression signature that, in turn, renders these cells more sensitive to IFNγ-induced apoptosis. Our findings reveal an age-related imbalance in the interaction between the immune and epithelial compartments in the colon, priming the system for excessive inflammatory responses and the emergence of a hypersensitive epithelial cell state thus derailing proper repair of the intestinal epithelium after injury.

## Main

Aging is characterized by a progressive decline in tissue function ^1^. This decline is associated with a chronically elevated inflammatory state, also termed inflammaging, of likely multifactorial origin, such as lifelong exposure to various antigens, accumulation of cellular and molecular damage, and changes in the microbiota ^2–4^. Aging further increases vulnerability to and incidence of pathogenic infections, exacerbating the chronic inflammatory state ^3, 5–9^. These age-related changes are particularly prevalent in barrier epithelia like the intestinal epithelium. To maintain tissue function despite frequent environmental challenges, the intestinal epithelium is highly regenerative, relying on intestinal stem cells (ISCs) and transit-amplifying cells (TAs) to restore homeostasis after damage. Regenerative capacity depends on tightly controlled interactions of these cells with their niche and the immune compartment in the lamina propria ^10–14^. Accordingly, epithelial-immune interactions are crucial for proper regeneration during infection-induced injury^13–19^.

Inflammatory challenges to the healthy adult intestine, such as infection with pathogens, activate the innate and adaptive immune systems ^15, 20, 21^. Epithelial cells secrete antimicrobial proteins that trigger the activation of macrophages and neutrophils, which combat the infection through cytokine production and phagocytosis of bacteria. Dendritic cells aid in this defense by directing lymphocytes to the infection site, and CD4^+^ T cells, B cells, and innate lymphoid cells (ILCs) are necessary for pathogen clearance through cytokine (IFNγ, IL17A, IL22) and antibody (IgG) production ^15, 20, 21^. During this response, cytokines such as IL22 promote crypt cell proliferation and regeneration ^22–26^. The healthy adult epithelium has also been shown to respond to IFNγ, secreted by lymphocytes, which induces transient transcriptional reprogramming to a fetal-like state, repurposing aspects of fetal development to regenerate successfully ^17–19^. This IFNγ-induced fetal-like reprogramming has been well-documented as a critical regenerative mechanism in response to injury and infection, both in the small and large intestine, where it allows for rapid epithelial restoration following damage. The crosstalk between the epithelium and immune cells in adulthood has been studied ^27–29^, but whether and how this crosstalk is affected by aging and impacts age-related regenerative dysfunction remain to be characterized.

The aging small intestine of the mouse undergoes relatively minor epithelial changes, such as decreased ISCs and impaired proliferation and differentiation ^30–33^. These changes result in a diminished regenerative response to various chemical-induced injuries, including 5-fluorouracil (5-FU), irradiation, and dextran sulfate sodium (DSS) ^31, 32, 34–37^. It has recently been discovered that excessive IFNγ production increases antigen-presenting pathway (APP) genes and biases towards the secretory lineage in the proximal small intestinal epithelium, negatively affecting regeneration ^28, 29^. Compared to the proximal gut, the distal gut is more commonly affected by intestinal tract disorders, such as inflammatory bowel disease, diverticulitis, and cancer ^38–41^. The higher bacterial load naturally present in the distal gut along with slower transit time and anaerobic conditions, increases the risk for infections in this region of the intestine ^42, 43^. Despite its clinical relevance, research on the effects of inflammaging in the distal gut is limited.

Here, we surveyed the immune and epithelial compartments of young and aged mouse colon in homeostasis and during infection-induced inflammation. During homeostasis, we found evidence suggesting an immune-primed state in the aged colon. Notably, this was not accompanied by a significant increase in baseline epithelial inflammation, contrasting with previous reports for the small intestine ^28, 29^. However, we observed a stark difference in inflammatory dynamics between old and young guts in response to *Citrobacter rodentium* infection, which elicits a robust innate and adaptive inflammatory response in the colon. After infection, we observed a significant age-related decline in *Lgr5*^+^ ISCs, associated with impaired regeneration, as well as with more rapid and higher induction of inflammation. We demonstrate that these changes are associated with two concomitant phenotypes uniquely present in the old gut: (1) elevated IFNγ response signature in epithelial cells and (2) enhanced induction of fetal-like genes associated with epithelial regeneration processes and cellular de-differentiation. Using organoid models, we show that cells in a fetal-like state, when exposed to sustained IFNγ signals, undergo programmed cell death, suggesting that an earlier and overactive pro-regenerative fetal signature in the epithelium of old injured mice may in fact make them more susceptible to inflammation-induced damage. Overall, our study suggests that elevated IFNγ exposure hinders intestinal repair in aged mice by maintaining a hypersensitive, fetal-like epithelial state.

## Results

### Transcriptional profiling reveals immune-specific inflammatory signatures in the aged mouse colon

To assess age-related cell type composition and transcriptional changes in the intestinal colonic epithelium, we performed 10X Genomics scRNAseq on sorted epithelial cells from the colon of young (2-5 months) and old (24-32 months) mice. Epithelial cells from the colon were isolated and the sequencing dataset was integrated with single-cell data from a previous publication (see Materials and Methods) ^44^ to create a more robust representation of aging colonic epithelium cell phenotype. For the analysis of the immune compartment, we utilized the single-cell data from the same previous publication ^44^. Using an unsupervised clustering approach after data integration, we defined distinct cellular populations in both epithelial and immune compartments (Figure 1A, D) that were annotated via the expression of key marker genes (Figure S1)

**Figure 1.**
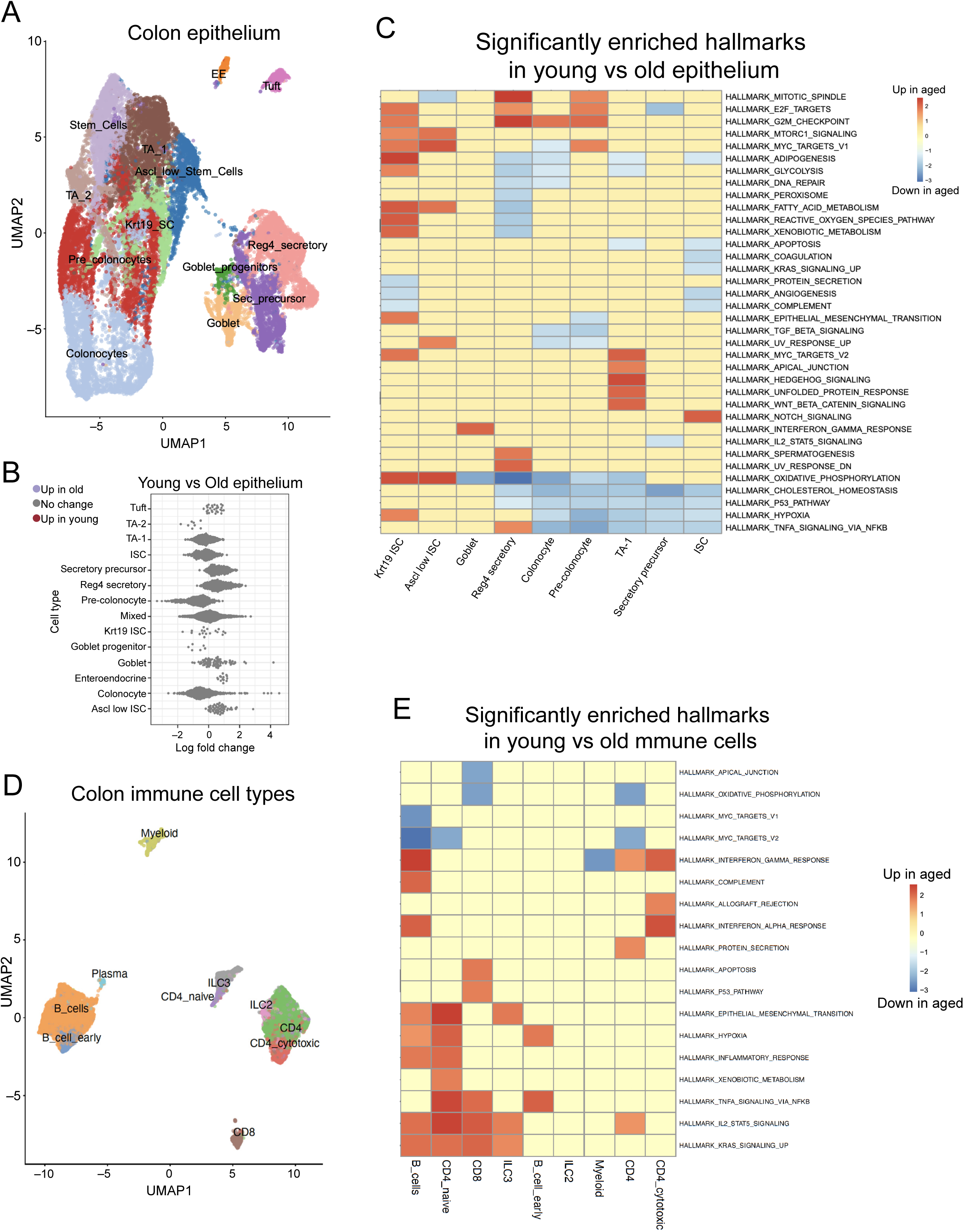
Transcriptional profiling reveals immune-specific inflammatory signatures in the aged mouse colon. (A) UMAP plots of young and aged mouse colon epithelial cell types. (B) Cell proportion analysis of epithelial cell types in young and aged mouse colon. Colored dots represent significantly differentially abundant cell neighborhoods with a spatially adjusted FDR of < 0.1. (C) Significantly enriched Hallmarks in the epithelial cell types of the mouse colon. Shown are the NES values calculated by the fgsea program. (D) UMAP plots of young and aged mouse colon immune cell types. (E) Significantly enriched Hallmarks in the immune cell types of the mouse colon. Shown are the NES values calculated by the fgsea program. (E) Significantly enriched Hallmarks in the epithelial cell types of the mouse ileum and colon.

To look for differences in cell type proportions between young and old while accounting for differences between integrated studies, we used the Milo framework ^45^, a linear model framework for estimating changes in cell type “neighborhood” proportions while accounting for differences in batch or experiment-specific effects We observed no statistically significant changes in epithelial cell type composition (Figure 1B), though we did note a sub-significant increase in the Ascl2-low stem cell population previously noted by Sirvinskas *et al.* ^44^, suggestive of a general reduction in stem cell activity with age (Figure S2A, B).

While epithelial cell type composition remained relatively stable with age overall, we observed modest but consistent changes in gene expression patterns (Figure S3A). To understand these changes, we performed a gene set enrichment analysis (GSEA) for differential expression within each cell type using the Hallmark gene set ^46, 47^. Interestingly, the aged colon epithelial cells did not exhibit a significant enrichment for IFNγ and IFNα responses or other hallmarks related to inflammation (Figure 1E). Instead, colonic epithelium cells upregulated cell cycle-related genes and hedgehog signaling while downregulating pathways involved in oxidative phosphorylation, cholesterol, and TNFα signaling (Figure 1C).

### Age-related expansion of pro-inflammatory immune cell populations in the colon

GSEA hallmark analysis of immune cells gene expression patterns from reference^44^ revealed that almost all resident immune cell types (CD8^+^ T cells, ILC2s, ILC3s, B cells, and CD4^+^ T cells), with the exception of myeloid cells, exhibited significant age-related increases in expression of inflammatory pathways, including IFNγ, IFNα, TNFα, and IL2 signaling (Figure 1E). This is in contrast to our observation in the epithelial compartment and suggested potential compartment-specific inflammatory signature emerging in the aging intestine. To test this idea and directly investigate whether the transcriptional changes we observed in the immune scRNAseq analysis were accompanied by shifts in immune cell composition, we isolated cells from colon lamina propria of young (2-3 months) and old (20-22 months) mice and performed flow cytometric analysis. We employed a comprehensive gating strategy to identify and quantify major immune cell populations and their subsets within the CD45^+^ compartment (Figure 2A).

**Figure 2.**
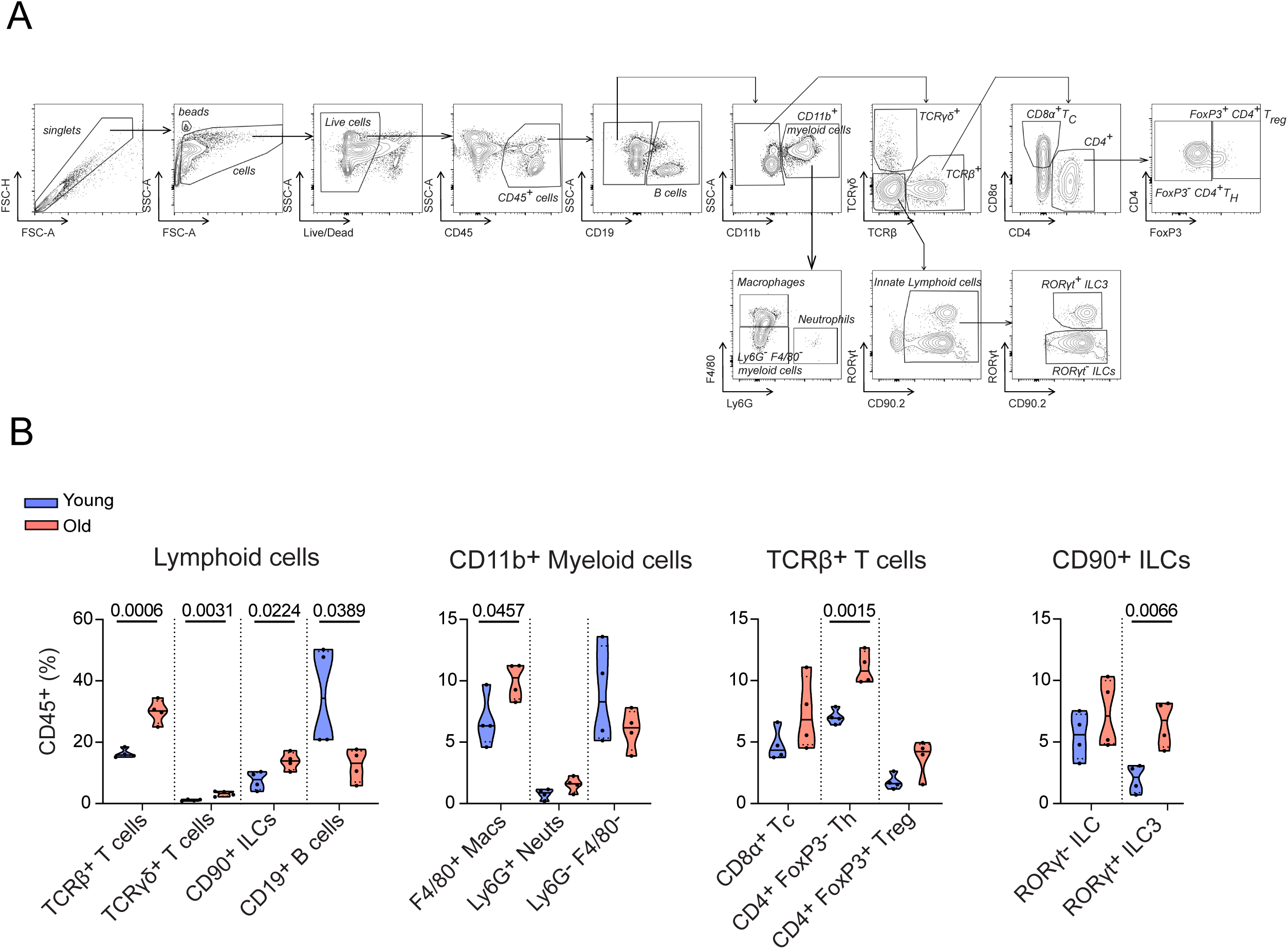
Age-related expansion of pro-inflammatory immune cell populations in the colon (A) Flow cytometry strategy followed for identification of immune cells in the colon lamina propria. (B) Cell proportions of colon immune cells in young and aged mice showing a shift towards pro-inflammatory cells in old animals. Data are presented as mean and SD (4 biological samples). Statistical comparisons were performed using unpaired t-test (*P*≤0.05).

Flow cytometric analysis revealed significant age-related changes in immune cell composition in the colon lamina propria (Figure 2B). We observed a substantial increase in total T cell numbers (both TCRβ and TCRγδ subsets), and a marked decrease in B cell numbers with age (Figure 2B). More detailed analysis of T cells subpopulations showed that CD4^+^ T helper cells were significantly increased, while CD8^+^ T cells and FoxP3^+^ regulatory T cells showed increasing trends that did not reach statistical significance. These T cell alterations align with enhanced T cell inflammatory signatures observed in our transcriptional analysis (Figure 1E).

Additionally, we detected a significant increase in ILC3s, which are known producers of pro-inflammatory cytokines in intestinal tissue. The aged colon also harbored more F4/80^+^ macrophages compared to young controls. Together, these compositional changes reveal a fundamental shift in the immune landscape of the aged colon, characterized by increased prevalence of cell types associated with inflammatory responses.

### Old mice are more vulnerable to enteropathogenic infection, in part due to impaired epithelial regeneration

Our survey of the aging distal gut reveals changes in immune cell transcriptomes and composition in old colons consistent with inflammaging, but with little evidence of changes in transcriptomes or cellular composition in the epithelium at homeostasis. We wondered whether the changes in the immune compartment may influence the epithelial response to infection, and infected mice with *C. rodentium*, a gram-negative murine bacterium that infects the distal gut, primarily colonic enterocytes, leading to acute colitis, and activation of both an innate and adaptive immune response ^21, 48–50^. Young (2-3 months old) and old (20-22 months) mice received 2 x 10^9^ CFU of *C. rodentium*, and their body weight and survival were monitored for three weeks (Figure 3A). Young animals exhibited short-term weight loss starting at around day (D) 8 post-infection, with full recovery by D17, while old animals exhibited significant body weight loss as early as D2 after infection and continued to decline without recovering (Figure 3B). By D16 all aged animals had succumbed, while almost all young animals survived (Figure 3C).

**Figure 3.**
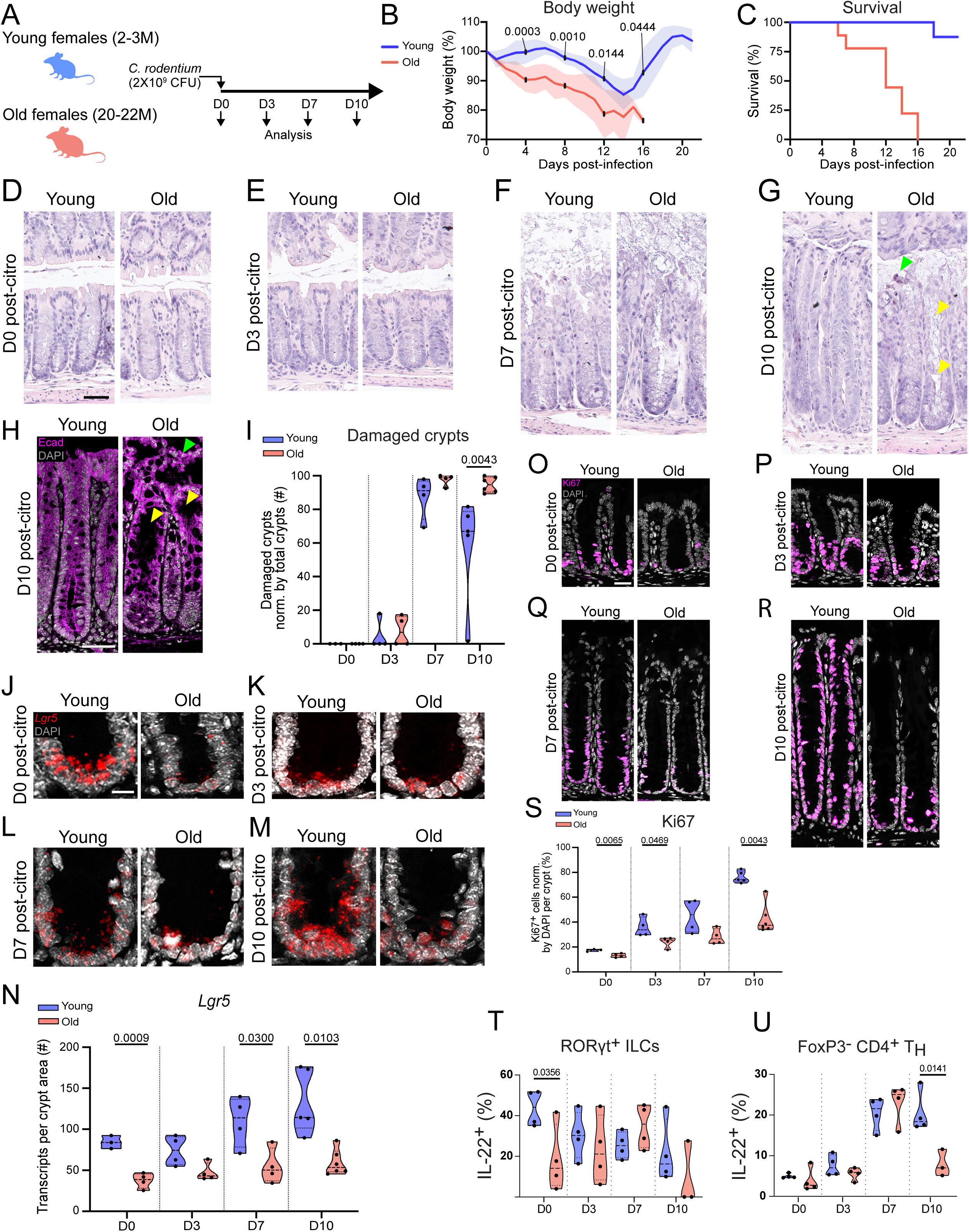
Aged colons exhibit increased epithelial damage and reduced ISCs and proliferation after infection. (A) Schematic of the experimental plan to assess the effect of aging in intestinal regeneration during infection. (B) Body weight loss during infection in young vs aged animals. Data are presented as mean and SD (8 animals per age group were enrolled in the study). Statistical comparisons were performed using unpaired t-test with Welch’s correction (*P*≤0.05). (C) Survival during infection in young vs aged animals. Data are presented as mean and SD (8 animals per age group were enrolled in the study). (D-G) HE staining of young and aged colons in homeostasis and during infection (D3, 7, and 10 post-*C. rodentium*) exhibit age-related epithelial damage during infection. Green arrowhead depicts detached inter-crypt epithelium shedding into the intestinal lumen and yellow arrowheads depict loss of epithelium and columnar morphology within crypts. Scale bar: 40 μm. (H) E-cadherin staining of young and aged colons on D10 post-*C. rodentium*. Green arrowhead depicts detached inter-crypt epithelium shedding into the intestinal lumen and yellow arrowheads depict loss of epithelium and columnar morphology within crypts. Scale bar: 40 μm. (I) Quantification of damaged crypts in young and aged colons from the HE stains (D-G). Data are presented as mean and SD (3-6 animals per age and timepoint were used for quantification). Statistical comparisons were performed using unpaired t-test with Welch’s correction (*P*≤0.05). (J-M) RNAscope for *Lgr5* transcripts in young vs aged colonic crypts during homeostasis and infection indicating reduced stem cell number or function in older animals (D0, 3, 7, and 10 post-citrobacter). Scale bar: 10 μm. (N) Quantification of *Lgr5* transcripts in young vs aged colonic crypts during homeostasis and infection (D0, 3, 7, and 10 post-*C. rodentium*). Data are presented as mean and SD (3-6 animals per age and timepoint were used for quantification). Statistical comparisons were performed using unpaired t-test with Welch’s correction (*P*≤0.05). (O-R) Ki67 staining for assessment of proliferation in the crypts of young and aged colons during homeostasis and infection (D0, 3, 7, and 10 post-*C. rodentium*). Scale bar 40 μm. (S) Quantification of Ki67^+^ proliferative cells in colon crypts of young vs aged animals during homeostasis and infection (D0, 3, 7, and 10 post-*C. rodentium*). Data are presented as mean and SD (3-6 animals per age and timepoint were used for quantification). Statistical comparisons were performed using unpaired t-test with Welch’s correction (*P*≤0.05). (T) Quantification of IL22^+^ RORγt^+^ ILCs in colon lamina propria of young vs aged animals during homeostasis and infection (D0, 3, 7, and 10 post-*C. rodentium*). Data are presented as mean and SD (3-4 animals per age and timepoint were used for quantification). Statistical comparisons were performed using unpaired t-test with Welch’s correction (*P*≤0.05). (U) Quantification of IL22^+^ of FoxP3- CD4^+^ T helper cells in colon lamina propria of young vs aged animals during homeostasis and infection (D0, 3, 7, and 10 post-*C. rodentium*). Data are presented as mean and SD (3-4 animals per age and timepoint were used for quantification). Statistical comparisons were performed using unpaired t-test with Welch’s correction (*P*≤0.05).

The heightened susceptibility to infection in old mice was accompanied by evident changes in the colonic epithelium. Histological analysis by HE staining and E-Cadherin immunostaining, show that the overall structure of colonic crypts in both young and old animals was similar and undisturbed before and on D3 after infection. By D7 post-infection, both young and old animals exhibited a significant increase in *C. rodentium*-induced epithelial damage, featuring the effacement of colonocytes (Figures 3D-F, I). By D10, young animals showed evidence of epithelial recovery, with about 60% of crypts in the distal 0.3 cm of the colon identified as damaged, whereas aged animals maintained a high proportion of damaged crypts (about 95%; Figures 3G, I).

To gain more granular insight into *C. rodentium*-induced epithelial damage in young and old animals, we performed immunostaining for epithelial polarity and BM markers as proxies for epithelial barrier disruption. The epithelial barrier is dependent on epithelial cell polarity ^50^, as well as on the basement membrane (BM). The BM acts as a physical barrier and separates the epithelium from the underlying mesenchyme and the lamina propria, where a plethora of immune cells reside ^51^.

To assess epithelial polarity, we stained for Ezrin, a protein localized in the apical membrane of epithelial cells, and required for the maintenance of intestinal epithelium homeostasis ^52^. Prior to infection and on D3 after infection, both young and old animals exhibited comparable levels of apical Ezrin protein expression (Figure S4 A-E). By D7, Ezrin levels in both young and old groups were reduced, suggesting that both groups had undergone epithelial damage. Notably, older animals displayed a more severe loss of Ezrin than young animals. Only 14% of old epithelium had intact apical Ezrin expression in comparison to 38% of Ezrin-positive apical expression in young. These differences were retained at D10 post-infection.

We further stained for laminin, one of the main ECM components of the BM. Interestingly, the area of the epithelium exhibiting laminin expression was reduced in aged epithelium compared to young at baseline prior to infection and at D3 after infection (Figure S4 F-G, J). While both age groups had a decline in laminin by D7, this decline stabilized in the young animals by D10, while old mice showed a further decrease at D10 compared to the young (Figure S4 H-J).

Previous studies in young colons have demonstrated that the main epithelial cell types responsible for successful regeneration after *C. rodentium* infection are ISCs and TAs ^53–56^. To better understand whether the increased epithelial damage and inability to regenerate in the aged group was due to reduced ISCs and/or TAs proliferation, we next focused on those cell types and their dynamics in young and old colons prior to and during infection. We first performed RNAscope for *Lgr5* transcripts, a widely used marker for ISCs. Prior to infection, aged distal colons showed a decrease in *Lgr5* transcripts per crypt (Figures 3J, N), suggesting a potential age-related decrease of ISCs in accordance with previous studies ^53–56^. By D3 post-infection, crypts of young animals showed a gradual elevation in *Lgr5* transcript numbers (Figures 3K-N), reflecting the induction of a regenerative response ^53–56^. In contrast, during the peak of the regenerative phase (D7 to D10), old crypts showed lower *Lgr5* expression (Figures 3L-N), and such expression was restricted to crypt bottoms. This was consistent with lower proliferation in crypt cells of aged mice compared with young at almost all time points post-infection, as assessed by Ki67 staining (Figures 3O-S).

The immune compartment of the colon also showed significant age-related changes in the response to infection. During *C. rodentium* infection in young mice, ILCs and T helper cells produce IL22, a cytokine required for further increasing crypt proliferation and inducing crypt hyperplasia, a crucial step for successful epithelium regeneration ^22–26, 57^. In colons of young animals, CD4 T helper cells progressively increased their expression of IL-22, whereas RORγt^+^ ILC3s had a high initial expression of IL-22, which gradually decreased throughout the infection (Figure 3T, U; Figure S5A). Interestingly, IL-22 production from CD4^+^ T helper cells in aged colons differed at the D10 time point, where the production dropped dramatically (Figure 3U; Figure S5A). Furthermore, aged ILC3s already exhibited lower levels of IL-22 production at baseline and did not notably change during the infection (Figure 3T; Figure S5A). These results indicate that older mouse colons are more sensitive to damage caused by *C. rodentium* compared to younger mice, likely attributable to a diminished ability to regenerate, which is characterized by a decrease in proliferation and ISCs. Furthermore, the observed decrease in IL-22^+^/CD4^+^ T helper cells and the relatively reduced production of IL-22 in aging animals supports the impaired epithelial regeneration after *C. rodentium* infection, which may contribute to the observed weight loss and mortality.

### Age-associated adaptive immune response to bacterial infection is characterized by a premature onset and elevated production of IFNγ

To better define changes in biological mechanisms taking place in young vs aged groups at the different times of infection, we performed bulk RNA-seq and assessed transcriptomic differences between young and old colons (including both epithelial and immune compartments) across the time course of infection (Figure 4A). We identified genes that changed expression significantly as a function of age (either directly or in an interaction with time point in a linear model), clustered these genes by their pattern of expression over the time course using the degPatterns package^58^ (Figure S6), and performed gene set enrichment analysis on each resulting cluster to better understand the biological processes influenced by genes on each cluster (Figure S7).

**Figure 4.**
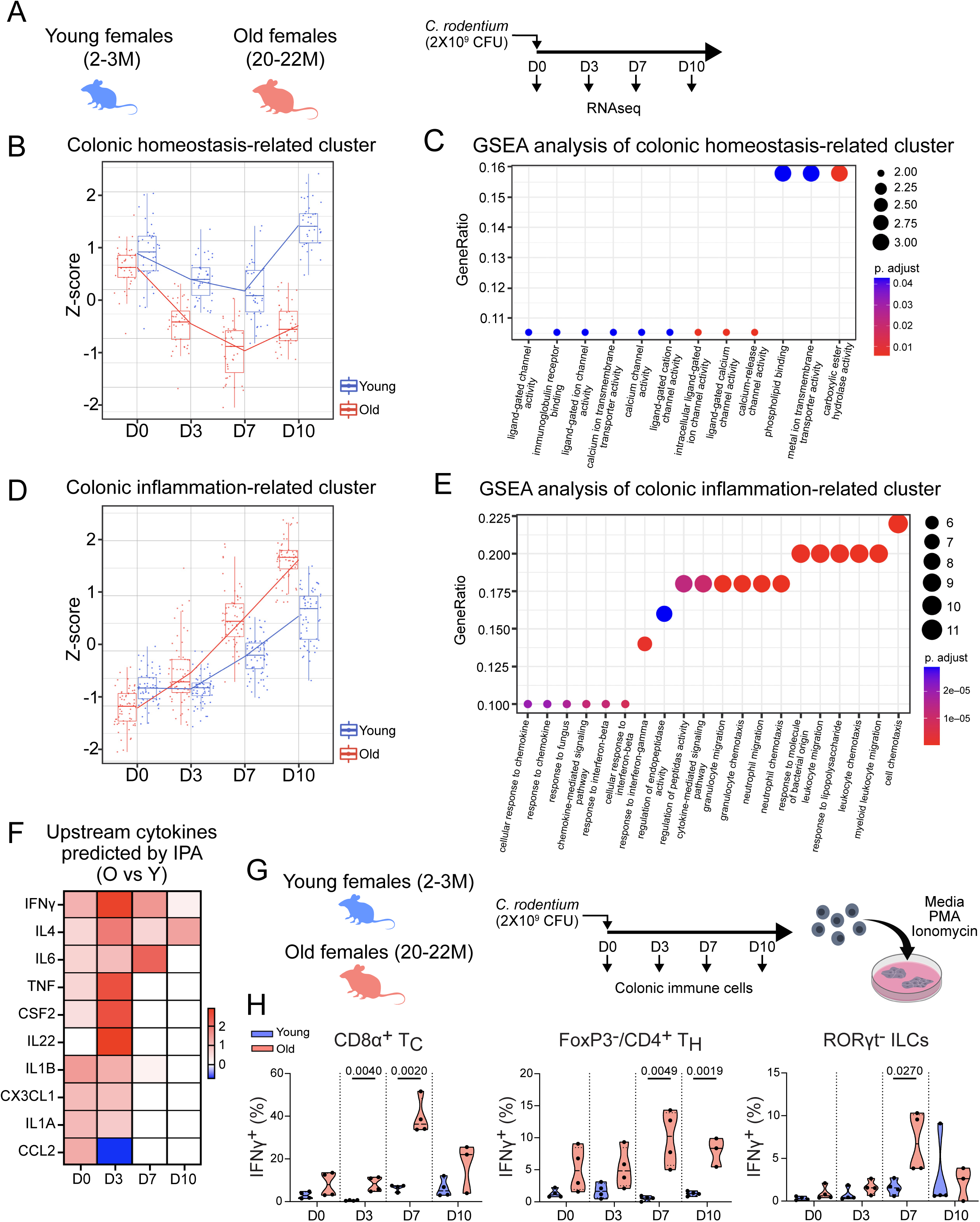
Aged-associated immune response to bacterial infection is characterized by a premature onset, elevated expression, and prolonged production of IFNγ. (A) Schematic of the experimental plan to assess the colonic transcriptome in young and aged animals during infection. (B) Z-score of a cluster containing DE genes related to overall colonic function of young and aged colons during homeostasis and infection. (C) GSEA analysis of cluster from (C) identifying hallmarks related to colonic function. (D) Z-score of a cluster containing DE genes related to inflammation of young and aged colons during homeostasis and infection. (E) GSEA analysis of cluster from (E) identifying hallmarks related to colonic function. (F) IPA-based prediction of upstream cytokines in aged vs young colon transcriptome during homeostasis and infection. (G) Schematic of the experimental plan to assess IFNγ production in young and aged colonic immune cells. (H) Percentage of IFNγ^+^ CD8a^+^ T, FoxP3^-^/CD4^+^ T, and RORγt^-^ ILC cells in young and aged colons during homeostasis and infection (D0, 3, 7, and 10 post-*C. rodentium*). Data are presented as mean and SD (3-4 animals per age and timepoint were used for quantification). Statistical comparisons were performed using unpaired t-test with Welch’s correction (*P*≤0.05).

Analysis of age-dependent differential gene expression revealed two prominent themes. The first theme, encompassing Clusters 5, 6, and 7 (Figures S6, S7), involved genes crucial for colonocyte homeostasis. These included transporters and enzymes essential for water and nutrient absorption (e.g., *Slc36a2*, *Slc30a10*), lipid metabolism (e.g., *Cyp2c55*), and ion channel function (e.g., *Trpa1*, *Car1*) ^59^ (Cluster 6 detailed in Figures 4B-C; Figure S8). In young mice, expression within Cluster 6, used as a representative example, transiently decreased until D7 post-infection but was fully restored by D10 (Figure 4B). This transcriptional recovery mirrored the successful epithelial regeneration observed histologically in young animals by D10 (Figures 3G-I, S4). In stark contrast, aged colons exhibited a more pronounced and persistent downregulation of these homeostatic genes, with expression failing to recover by D10 (Figure 4B). This lack of transcriptional restoration aligns with the persistent epithelial damage we observed in aged colons at D10 (Figure 3I). Furthermore, the sustained reduction in colonocyte marker expression in aged mice from D7 onwards (Figure S8) suggests a failure to replenish these cells post-infection, reinforcing the conclusion that the aged epithelium exhibits impaired repair capacity.

The second major theme, represented by Clusters 1, 9, 15, and 30 (Figures S6, S7), comprised genes strongly associated with inflammation (e.g., *Ccl3*, *Timp1*, *Arg1*, *Ccl2*, *Ccl7*, *S100a8*, *S100a9*). Pathway analysis of these clusters (Cluster 1 detailed in Figures 4D, E; Figure S9) showed enrichment for hallmark pathways related to innate immune cell chemotaxis and responses to IFNγ and IFNβ cytokines (Figure 4E). Compared to young animals, aged mice displayed a markedly exacerbated upregulation of genes in Cluster 1 throughout the infection course (Figure 4E), indicating an intensified inflammatory response specifically triggered by the infection. Collectively, these data demonstrate that aged colons mount an excessive and prolonged inflammatory response to *C. rodentium*. This heightened inflammation correlates temporally with the earlier onset, increased severity, and unresolved epithelial damage observed in aged mice (Figures 3, S4), strongly suggesting that this dysregulated inflammatory environment contributes to the impaired regenerative capacity of the aged colonic epithelium.

As an additional method to analyze and predict upstream regulators of the observed gene expression changes throughout the time course of infection, we utilized Ingenuity Pathway Analysis (IPA) ^60^. Since many genes affected by aging were related to immune cell recruitment and cytokine response, we focused our analysis on predicting potential upstream cytokine regulators. IFNγ was predicted to be the top upstream cytokine, which increased with age at all time points (Figure 4F). To validate these predictions, we performed FACS analysis on lamina propria immune cells throughout the infection time course. We confirmed elevated IFNγ protein expression in CD8⍺^+^ T cells, CD4^+^ T helper cells, and RORγt^−^ ILCs after PMA and ionomycin stimulation in old animals (Figure 4G; Figure S10A). This finding is consistent with the increased numbers of T cells and ILC3s we observed in aged colons at homeostasis (Figure 2B), which likely contribute to the elevated IFNγ production during infection. This also complements our earlier observation of altered cytokine responses in aged animals, where we saw diminished IL-22 production in ILC3s and CD4+ T cells (Figure 3T-U), suggesting an imbalance in the inflammatory-regenerative axis. Notably, old animals already exhibited a non-significant increase in IFNγ production in these cell types at baseline, supporting our hypothesis that immune cells in aged colons are primed toward an inflammatory response, as indicated by our homeostatic transcriptional data (Figure 1E).

### Aging enhances epithelial sensitivity to IFNγ

Previous studies have shown that IFNγ can induce damage to the proximal intestinal epithelium in both young adult mice and human organoids ^13, 14, 28, 61^. Furthermore, a recent study has demonstrated that treating mouse duodenum organoids with IFNγ recapitulates the reduction in LGR5^+^ ISCs, increase in the secretory cell lineage, and induction of the antigen-presenting pathway observed in the aging intestinal epithelium ^34^. These data and our findings led us to hypothesize that the aging colon is more susceptible to IFNγ-induced damage. To test this hypothesis, we established colonic organoids from young and aged mice, exposed them to PBS or IFNγ, and monitored their survival over a 60h period (Figure 5A). Consistent with previous reports on the effects of aging on colonic organoid formation ^24^, aged crypts showed significantly reduced organoid formation capacity, and these organoids exhibited elevated cleaved caspase 3/7 staining. IFNγ treatment resulted in reduced survival and increased cell death in young colonoids (Figures 5A, B; S11A-B). A similar IFNγ-mediated phenotype has been described in the small intestine ^28^. Interestingly, organoids from old colons were significantly more sensitive to IFNγ than organoids from young colons, showing elevated cleaved caspase 3/7 staining and organoid death after IFNγ treatment (Figures 5A, B; S11A-B). This increased sensitivity to IFNγ may explain the inability of old intestines to recover from *C. rodentium* infection.

**Figure 5.**
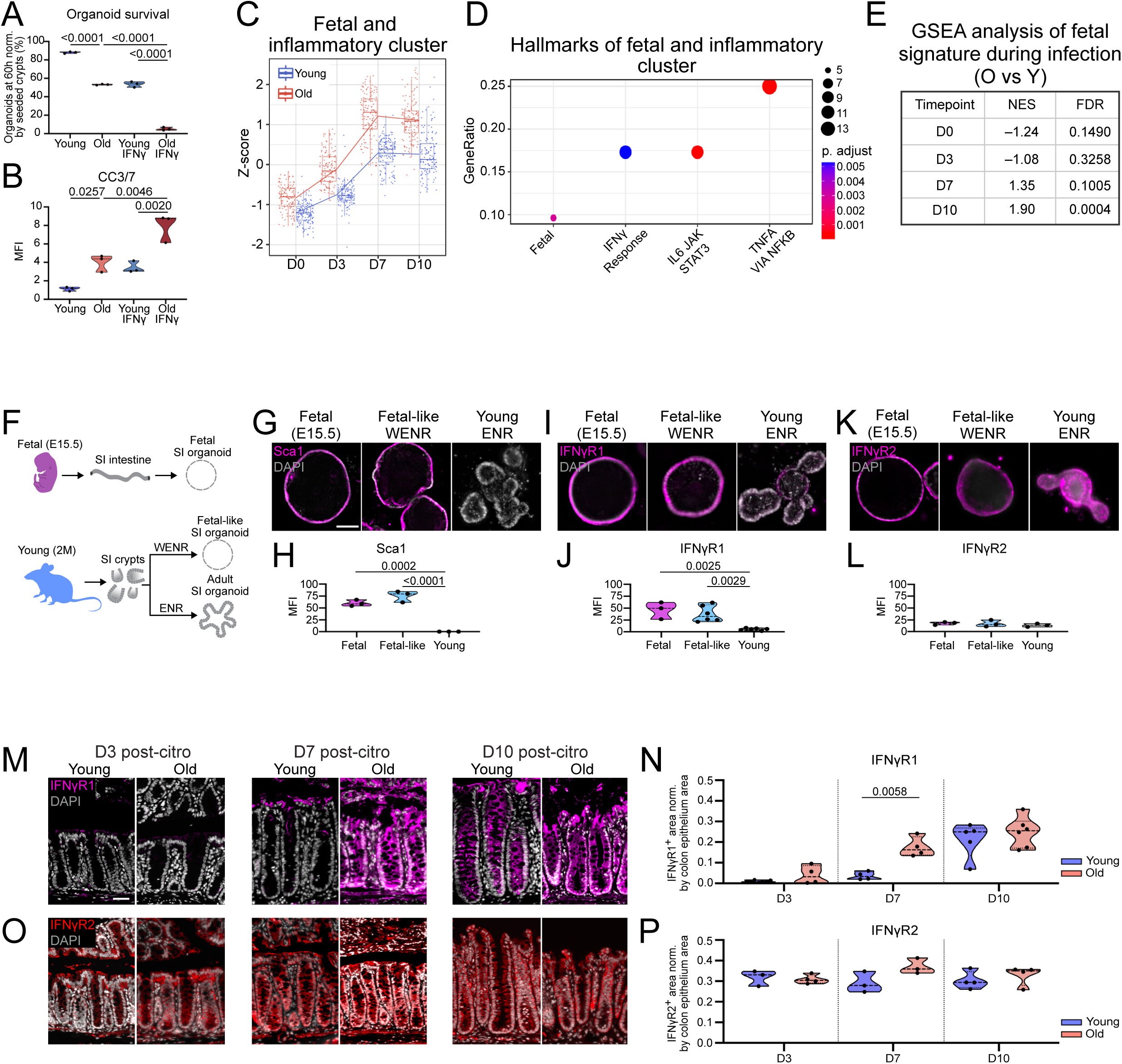
Elevated IFNγ exposure in aged epithelium induces sustained fetal-like reprogramming, increasing cell sensitivity to IFNγ-mediated damage. (A) Quantification of the percentage of organoid survival in aging and after IFNγ treatment. Data are presented as mean and SD (3 biological samples per age were used for quantification). Statistical comparisons were performed using a one-way ANOVA with Tukey’s test (*P*≤0.05). (B) Quantification of apoptotic CC3/7^+^ organoids in aging and after IFNγ treatment. Data are presented as mean and SD (3 biological samples per age were used for quantification). Statistical comparisons were performed using a one-way ANOVA with Tukey’s test (*P*≤0.05). (C) Z-score of a fetal and inflammatory gene set cluster in young and aged colons during homeostasis and infection. (D) GSEA analysis of the cluster from (C) identifying hallmarks related to fetal and inflammatory pathways. Dot size encodes gene count. (E) GSEA analysis of fetal signature in pairwise DE comparisons between young and old in each time point (D0, 3, 7, and 10 post-*C. rodentium*). (F) Schematic of the experimental plan to generate fetal (E15.5), fetal-like (WENR cultured), and adult (ENR cultured) intestinal organoids. (G) Sca1^+^ staining of fetal, fetal-like (WENR), and adult (ENR) intestinal organoids. Scale bar: 30 μm. (H) Quantification of (G) Sca1^+^ mean fluorescent intensity (MFI) in organoids from fetal, fetal-like (WENR), and adult (ENR) intestinal organoids. Data are presented as mean and SD (3 biological samples per group were used for quantification). Statistical comparisons were performed using a one-way ANOVA with Tukey’s test (*P*≤0.05). (I) IFNγR1^+^ staining of fetal, fetal-like (WENR), and adult (ENR) intestinal organoids. (J) Quantification of (I) IFNγR1^+^ MFI in fetal, fetal-like (WENR), and adult (ENR) intestinal organoids. Data are presented as mean and SD (3-6 biological samples per group were used for quantification). Statistical comparisons were performed using a one-way ANOVA with Tukey’s test (*P*≤0.05). (K) IFNγR2^+^ staining of fetal, fetal-like (WENR), and adult (ENR) intestinal organoids. (L) Quantification of (K) IFNγR2^+^ MFI in fetal, fetal-like (WENR), and adult (ENR) intestinal organoids. Data are presented as mean and SD (3-6 biological samples per group were used for quantification). Statistical comparisons were performed using a one-way ANOVA with Tukey’s test (*P*≤0.05). (M) IFNγR1^+^ staining of young and aged colons during infection (D3, 7, and 10 post-*C. rodentium*). Scale bar: 40 μm. (N) Quantification of IFNγR1^+^ colonic epithelium in young and aged colons during infection (D3, 7, and 10 post-*C. rodentium*). Data are presented as mean and SD (3-6 biological samples per age and group were used for quantification). Statistical comparisons were performed using unpaired t-test with Welch’s correction (*P*≤0.05). (O) IFNγR2^+^ staining of young and aged colons during infection (D3, 7, and 10 post-C*. rodentium*). (P) Quantification of IFNγR2^+^ colonic epithelium in young and aged colons during infection (D3, 7, and 10 post-*C. rodentium*). Data are presented as mean and SD (3-4 biological samples per age and group were used for quantification). Statistical comparisons were performed using unpaired t-test with Welch’s correction (*P*≤0.05).

### Prolonged IFNγ exposure drives an overactive and poorly resolved fetal-like reprogramming in aged epithelium, enhancing sensitivity to IFNγ-mediated death

We next sought to understand why the aged epithelium is more sensitive to IFNγ-mediated damage compared to the young by a deeper analysis of the degPattern gene clusters that were differentially affected by aging during infection (Figure 4). Of interest, one particular cluster was upregulated between old and young animals during our time course and enriched for IFNγ response as well as other inflammatory pathways such as IL6 and TNFα. Notably, this cluster was also enriched for a non-overlapping gene set defining a previously reported fetal gene signature (Figure 5C, table S1) ^17–19^. GSEA revealed a significant enrichment of this fetal signature at D10 after injury in the group of genes elevated in old compared to young samples (Figure 5D, E). This fetal signature represents a transcriptional program that is normally active during intestinal development, and reflects a state of increased epithelial cell plasticity and de-differentiation. Transient induction of this signature in adult tissues is associated with the capacity for rapid regeneration and repair. In the small intestine, it has been shown to be induced by IFNγ signaling and required for regeneration after parasitic infection, irradiation, or DTR-mediated ablation of LGR5^+^ ISCs ^17^.

We asked whether this fetal-like state would impact epithelial sensitivity to IFNγ signaling, and compared the IFNγ response of young organoids, fetal organoids, and ‘fetal-like’ organoids induced into a fetal-like state by exogenous WNT3A treatment ^18, 62^. We isolated and cultured fetal intestinal organoids (E15.5), and young (2 months) duodenum organoids, which were cultured in EGF, Noggin, and RSPO3 (ENR) media with or without the addition of WNT3A (Figure 5F). We used duodenum organoids for these experiments since their survival does not depend on exogenous WNT3A, contrary to colonoids, allowing us to experiment with two conditions (i.e., organoids in standard ENR media and organoids in ENR+WNT3A media). The morphological transition of budding organoids to spheres, and staining for the fetal marker Sca1, validated previous published work showing that the addition of WNT3A drives a fetal-like state in young organoids^18, 61^, while ENR-only cultured organoids had budding crypts and no expression of Sca1 (Figure 5G, H), as previously shown ^17, 63, 64^.

IFNγR1 and R2 receptors are required on the cell surface for IFNγ ligands to bind and initiate downstream activation of IFNγ signaling. To assess if differential availability of these receptors may explain the different sensitivity of organoids to IFNγ, we used immunohistochemistry to detect IFNγR1 and R2 in fetal, fetal-like, and young organoids (Figure 5I-L). While IFNγR2 was highly expressed across all organoids, IFNγR1 was expressed in both the fetal and fetal-like intestinal organoids, but its expression was lower in the young organoids (Figure 5I, J). A fetal state thus seems to be accompanied by increased expression of IFNγR1 (Figure 5I, J), potentially rendering it more sensitive to IFNγ-mediated damage and, eventually, cell death.

To validate our findings, we determined if the expression pattern of IFNγR1 and R2 observed *in vitro* is phenocopied in young and aged colons at D3, D7, and D10 post-infection (Figures 5M-P). Similar to our organoid data, on D3 post-infection, when a fetal signature was not yet induced, there is minimal staining of IFNγR1 in the young and aged epithelium (Figures 5M, N). However, by D7 post-infection, when a fetal signature started to be upregulated in old compared to young animals (Figure 5C-E), IFNγR1 expression was significantly increased in the aged epithelium (Figure 5M, N). Complementing our organoid studies, we found IFNγR2 to be expressed across the whole epithelium in young and aged colons without any significant differences prior to or after the fetal-like reprogramming of the epithelium during infection (Figure 5O, P).

Next, we tested whether a stable and prolonged fetal state would render the intestinal epithelium more vulnerable to sustained IFNγ exposure similar to the one seen in aging infected colons. In contrast to the young epithelium, fetal and fetal-like organoids were more sensitive to IFNγ-mediated death (Figure 6A-EQ-U). By D5 of culture with continuous exposure to IFNγ, fetal and fetal-like organoids accumulate dying cells and collapse.

**Figure 6.**
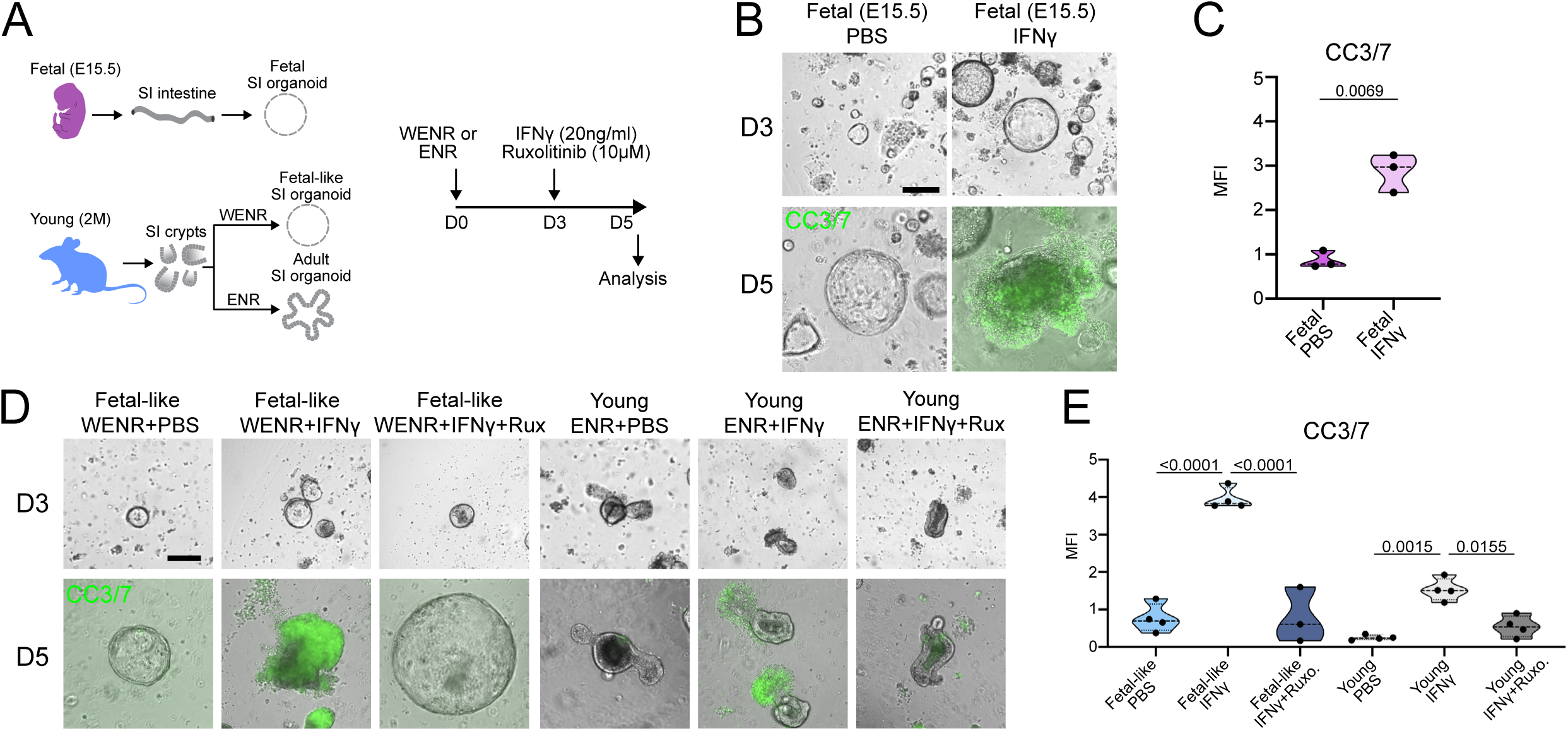
Inhibition of IFNγ signaling rescues the IFNγ-induced cell death in the fetal-like organoids (A) Schematic of the experimental plan to assess the effect of IFNγ and its downstream pathway inhibition via Ruxolitinib in fetal, fetal-like (WENR), and adult (ENR) intestinal organoids. (B) Representative images of CC3/7 staining of fetal intestinal organoids with or without IFNγ treatment. Scale bar: 30 μm. (C) Quantification of (R) CC3/7^+^ fetal intestinal organoids in control vs IFNγ-treated groups. Data are presented as mean and SD (3 biological samples per group were used for quantification). Statistical comparisons were performed using unpaired t-test with Welch’s correction (*P*≤0.05). (D) Representative images of CC3/7 staining of fetal-like (WENR), and adult (ENR) intestinal organoids under PBS (control), IFNγ, and IFNγ+Ruxolitinib conditions. Scale bar: 30 μm. (E) Quantification of (T) CC3/7^+^ fetal-like (WENR), and adult (ENR) intestinal organoids under treatment conditions. Data are presented as mean and SD (3-4 biological samples per group were used for quantification). Statistical comparisons were performed using a one-way ANOVA with Tukey’s test (*P*≤0.05).

IFNγ signaling proceeds via receptor engagement, JAK1/2 activation, and subsequent STAT1 phosphorylation, ultimately leading to the transcription of target genes involved in immune response and inflammation^42^. To determine if the IFNγ-JAK-STAT signaling axis mediates the vulnerability of the sustained fetal-like state to cell death induced by chronic exposure to IFNγ, we applied the JAK1/2 inhibitor Ruxolitinib concurrently with IFNγ treatment (Figure 6A). Indeed, JAK1/2 inhibition provided complete protection from IFNγ-induced damage. Ruxolitinib treatment caused a dramatic drop in cell death and preserved the characteristic spheroid structure typical of pro-regenerative fetal and fetal-like states, thereby fully rescuing the organoids from the detrimental effects of chronic exposure to IFNγ (Figure 6D, E).

## Discussion

A decline in resilience and an increase in chronic inflammation are hallmark characteristics of aging. In rapidly renewing organs such as the intestine, continuous and efficient stem cell-mediated regeneration is essential for maintaining tissue function both under homeostatic conditions and following injury. Understanding how aging affects this regenerative process, thereby contributing to a decline in resilience, remains a central question.

In this study, we have identified compositional and transcriptional changes in epithelial and immune cells of the aging colon. Our findings reveal a striking compartmentalization of aging effects, where immune cells exhibit clear inflammatory signatures while the epithelial compartment maintains relative transcriptional homeostasis. This suggests that the immune compartment, rather than the epithelium, is the primary driver of age-related inflammatory changes in the colon. This compartmentalization likely represents a state of “immune priming” in aged animals - where immune cells display pro-inflammatory transcriptional signatures and altered composition but remain below the threshold for full activation that would impact epithelial gene expression under homeostatic conditions. Such primed immune cells would be poised to mount an exaggerated response upon encountering a pathogenic trigger, explaining why dramatic inflammatory effects emerge only after challenge. This compartmentalization in the colon further represents a tissue-specific manifestation of ‘inflammaging’ that differs from what has been reported in the small intestine ^29, 34, 44^.

The resilience of the colonic epithelium to age-related changes when unchallenged, is consistent with recent observations ^44^, and may reflect its specialized adaptation to maintaining barrier integrity in an environment with higher bacterial loads and more challenging luminal conditions compared to the small intestine.

Our findings highlight an important distinction between inflammatory signatures and cytokine production in the aging intestine. While aged immune cells exhibit enhanced IFNγ response signatures at homeostasis, significant increases in actual IFNγ production are only observed following infection challenge, as demonstrated by our FACS data. This suggests that the aged immune system is primed toward an exaggerated inflammatory response but requires a trigger such as infection to manifest the full inflammatory phenotype. Our results identify IFNγ, most likely expressed by T cells and ILCs, as a key effector of inflammaging in the colon. While we identified lamina propria T cells and ILCs producing IFNγ, it is possible that additional sources of IFNγ, such as intraepithelial lymphocytes, contribute to the observed phenotype. We demonstrate that *C. rodentium* upregulation of IFNγ induces transcriptional changes that are not obvious at homeostasis, such as the upregulation of genes associated with the development of the fetal gut. This is consistent with previous work in the small intestine ^17–19^, and we propose that while the pro-regenerative fetal gene activation in epithelial cells after IFNγ exposure is beneficial in young, healthy guts, it becomes detrimental if unchecked. This unchecked activation impairs the return to homeostasis and damages the epithelial barrier in aged infected guts. Accordingly, previous studies have suggested that uncontrolled upregulation of IFNγ has a detrimental effect on ISCs and intestinal homeostasis in young/adults ^13, 65–67^.

Our data suggest that premature onset and elevated production of IFNγ signaling contribute to age-related impairments in intestinal regeneration. Nevertheless, additional molecular signals, such as IL-22 dysregulation, may also be influential. IL-22 is known to promote regeneration in both *in vivo* and *in vitro* settings by stimulating crypt cell proliferation ^24, 26, 49, 56^. During *C. rodentium* infection in young mice, Ki67 expression increases, indicating enhanced cell proliferation. However, this regenerative response is impaired in aged mice. Unlike young mice, aged mice exhibit a decrease in IL-22 production from FoxP3^-^ CD4^+^ T_H_ and RORγt^+^ ILCs, which may account for the diminished proliferative response observed in older mice. This impairment is further exacerbated by the already reduced presence of ISCs in the aged homeostatic colon, a phenomenon that mirrors the reduction in ISC numbers and function previously observed in the aged small intestine and colon ^30–32, 36, 43^. Investigating how the interplay of enhanced IFNγ signaling, diminished IL-22 response and compromised ISC proliferation influences the regenerative response presents a compelling area for future research. Furthermore, determining whether targeted modulation of these individual pathways can restore regenerative capacity will be of interest to explore.

During extensive intestinal damage where *Lgr5^+^* stem cells are insufficient to repair the epithelium, epithelial cells transition into a fetal-like state that leads to epithelial restitution ^17, 63, 68^. This fetal-like reversion is characterized by the upregulation of a genetic signature that is reminiscent of intestinal development. In the young small intestine, IFNγ and YAP have been identified as key inducers of this signature ^17, 63, 68^. Indeed, the role of IFNγ in driving this fetal-like reprogramming is a well-documented phenomenon, representing an evolutionarily conserved mechanism for epithelial repair following injury. Interestingly, the essential role of IFNγ and YAP in inducing fetal-like reversion to promote repair hinges on their transient activation, which enables cells to exit plasticity and differentiate back into a mature adult ISC state once the injury subsides. This reversion is critical for a balanced restoration of absorptive and secretory cell lineages ^68, 69^. In the aged colon, in turn, we find that *C. rodentium* infection leads to precocious and increased IFNγ upregulation compared to young colons and the aberrant induction of a fetal signature that includes elevated IFNγR1 expression. This causes elevated sensitivity to IFNγ, causing eventual cell death. While the pro-regenerative feedback loop between IFNγ and fetal gene activation can be beneficial in young animals, its imbalance in old animals impairs the return to homeostasis after infection, damaging the epithelial barrier and causing death (Figure 7). Our findings identify IFNγ signaling as a promising target for the treatment of age-related intestinal dysfunction, and indicate that timing and duration of IFNγ signaling are critical for successful intestinal regeneration.

**Figure 7.**
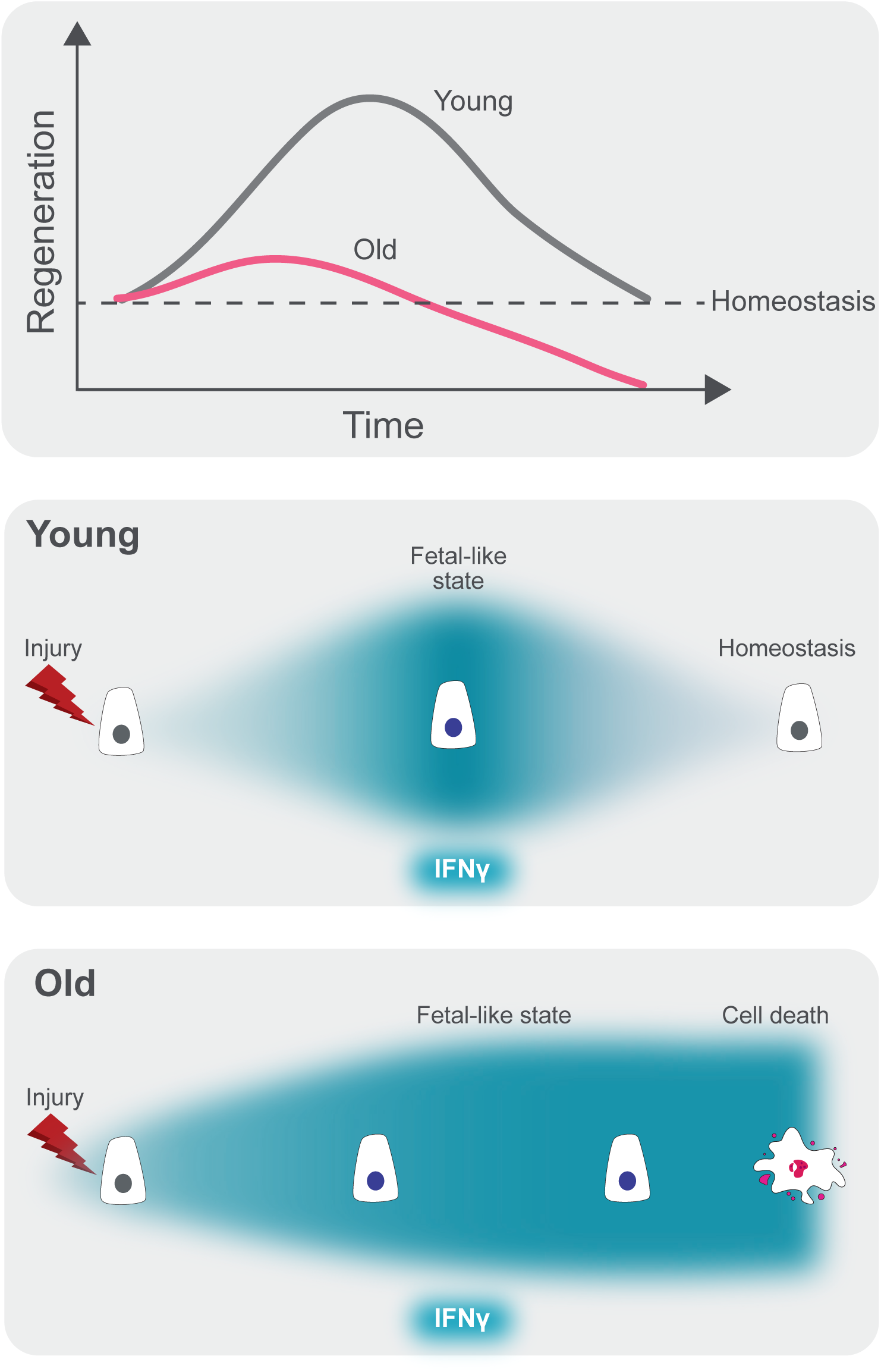
Model of colonic regeneration in aging. We have identified IFNγ as a key regulator of inflammaging. In young intestines, transient activation of IFNγ facilitates regeneration by upregulating genes that are commonly active during intestinal development. This brief reversion to a fetal-like state in healthy intestines creates an optimal condition—a “Goldilocks state”—characterized by the precise temporal activation of IFNγ and fetal genes. In contrast, prolonged upregulation of IFNγ in aged animals induces a persistent fetal-like state, which becomes susceptible to IFNγ-driven apoptosis and consequently impairs epithelial regeneration.

## Methods

### Mouse infections with *C. rodentium*

Mouse experiments were performed in accordance with the Institutional Animal Care and Use Committee (IACUC) and were approved by the Genentech IACUC. *C. rodentium* was purified on MacConkey agar, and one colony was inoculated in lysogeny broth (LB), and allowed to grow overnight to produce a stock solution. Young and aged mice were inoculated by oral gavage with 2 x 10^9 colony-forming units of *C. rodentium.* Body weights were monitored every day for the duration of the experiment and mice that lost 20% or more of their body weights were euthanized following IACUC standards.

### Colon alignment and atlas construction

Our single cell colon dataset of cells from old and young animals came from two sources: A collection of animals (two young at 4M old and three aged at 20, 21, and 28M old) processed in-house and the second from Sirvinskas et al. 2022 ^44^. The internal data were processed as described above with one exception – we set a hard threshold on mitochondrial percentage at 30% rather than using the outlier detection method. Samples were then clustered and given an initial annotation using a marker set (table S1) covering colonic and ileal cell types for subsequent processing steps. At this time, cells were split into epithelial and immune, with each set being processed individually.

We obtained the count matrices of Sirvinskas et al. 2022 as count matrix files from Abiosciences (ABS). For the immune and epithelial count matrices separately, we applied our standard approach of removing outliers based on MAD followed by clustering using the parameters above. While annotations from the publication were provided for the immune cells, the meta data for the epithelial cells did not contain cell type labels. We therefore projected our set of colonic and ileal markers onto the clustered epithelial data and annotated the cell types, updating the annotations as needed to match marker projections from the original publication.

To create the combined sample set we used in downstream analyses, we first applied a uniform filter that removed cells with > 15% mitochondrial content. We then split the data based on the initial, experiment-specific annotations into immune and epithelial subsets. To each subset independently, we applied the following procedure.

First, we transformed our count data using the logNormCounts.chan function from scran.chan using sample-specific size factors to downscale the size factors of all samples to match that of the least covered sample. We then modeled the per-gene variances across the dataset using the modelGeneVar.chan function, and selected the top 2000 most highly variable genes for downstream dimensionality reduction. This reduction was carried out using PCA as implemented in runPCA.chan with the batch method set to “block” (effectively regressing on experiment of origin as the batch variable prior to PCA for the epithelial cells and on sample for the immune cells). For the immune cells, we further adjusted the PCs by feeding the first 25 components to fastMNN ^70^ as implemented in scran.chan with k = 15. The resulting corrected PCs were then used for downstream analyses using the runAllDownstream function in scran.chan with the following parameters: tsne.perplexity = 30, umap.num.neighbors = 15, cluster.lmeans.k = 10, cluster.snn.method = “multilevel”, cluster.snn.num.neighbors = 10.

For the epithelial samples, we were not able to obtain an embedding with scran.chan that did not continue to separate the two experiments. We therefore made use of Seurat v4 ^71^. Specifically, we applied SCTransform to each Experiment while applying a regeneration on a log10 transformation of total counts as well as percent mitochondrial reads. We then selected the top 3000 variable features across the Experiments using the function SelectIntegrationFeatures followed by the anchor-based integration approach ^71, 72^. We then calculated the top 20 Principal Components using the 3000 most variable features followed by clustering and UMAP projection. Cluster identification was carried out using a resolution of 0.4. Annotations were then carried out by projection of our marker list (##supp figure) and examination of the cell identities from the original constituent studies.

### Milo analyses

To robustly evaluate changes in cell type proportions as a function of age, we made use of the Milo framework ^45^ which evaluates changes in condition-specific representation on a k-nearest neighbors graph that allows for the use of covariates such as batch/study effects. For all analyses, we used the default parameters with specific choices of “d” (the number of PCs to use for knn graph building) and “k” (the number of nearest neighbors) as follows: colonic epithelium, d = 25, k = 40; immune cells, d = 25, k = 30. For all of these analyses, which involved data from two experiments, we used the fastMNN adjusted PCs and included a term for experiment in the linear model. Statistical significance in the linear model contrasts was determined by having a spatially adjusted p-value of <= 0.1^76^.

### Pseudo-bulk differential expression analysis

To conduct robust analyses of differential expression that include the impact of biological variation among animals, we made use of a pseudo-bulk strategy and limma-voom ^73^. Briefly, total counts per gene were summed for each cell type and each sample to create a pseudo-bulk matrix. Any pseudo-bulk sample with fewer than 50 cells was removed from subsequent analyses. For each cell type, we then tested for an impact of age using a linear model that contained a term for Experiment using the limma package ^74^ as follows. First, sample specific quality weights were estimated using the voomWithQualityWeights function. We then fit the linear model to the log-transformed count data using lmFit and a call to eBayes to perform empirical Bayes shrinkage, sharing information across genes to stabilize variances.

### Gene expression clustering

To make sense of patterns of gene expression in old vs. young animals over the time course of the citrobacter infection, we first identified genes genes that changed significantly over the time course and/or showed a differential response to time using a linear model in limmaVoom (above) using a model that included a term for time and a time:Age interaction effect. The resulting nominally significant genes (p-value <= 0.05 and absolute log2FC >= 3) were clustered based on their expression patterns using degPatterns from the DEGreport package (v 1.32.0). These clusters were then annotated using enrichment of the GO and Hallmark gene sets within each cluster relative to all tested genes using “enrichGO” and “enrichr” (respectively) in the clusterProfiler^79^ package (v4.4.4.) with a Nejamini-Hochberg significance threshold of 0.05. We also evaluated the logFC and enrichment of genes in our fetal signature list (table S1) as a set using the cameraPR function from limma (v3.52.2).

### Intestinal immune cell isolation, sorting, and *in vitro* activation

Lamina propria cells were isolated from mouse colon using the mouse Lamina Propria Dissociation kit (Miltenyi). For restimulation, freshly isolated lamina propria cells were plated into a 96-well V-bottom plate (Corning) in complete RPMI media, and stimulated with 1x concentration of Cell Stimulation Cocktail (eBioscience) for 2 hours. Protein Transport Inhibitor Cocktail (eBioscience) was subsequently added for an additional 2 hours to allow for buildup of intracellular cytokines. After 4 hours of total restimulation, cells were washed with PBS, and stained with the Fixable Near-IR Dead cell stain kit (Invitrogen) to exclude dead cells. Cells were then washed with PBS, and stained with Fc receptor blockade using unlabeled anti-CD16/32 antibody (clone 2.4G2, BD Pharmingen), followed by surface staining with fluorophore-labeled antibody mixtures for various surface markers. For intracellular/intranuclear staining, cells were washed with FACS buffer, followed by overnight fixation using the eBioscience FoxP3/Transcription Factor Staining Buffer set (Invitrogen). After fixation, cells were washed with a permeabilization wash buffer, followed by addition of intra-cellular and intra-nuclear fluorophore-labeled antibody mixtures. Cells were subsequently washed, filtered through a 30-40 um 96-well filter plate (Acroprep), and resuspended in a FACS buffer. For absolute counts, 5 ul of CountBright Absolute Counting beads (Thermo Fisher) were added to each sample. Samples were then analyzed using a BD FACSymphony (BD Biosciences). The antibodies used for sorting the different immune cells as well as for quantifying different cytokines are summarized in table S2.

### Histology and immunofluorescence

Whole colon tissues were isolated, cut in 3 equal longitudinal lengths (proximal, mid, and distal regions), fixed in 4% paraformaldehyde (PFA) for 24 hours before being placed in 70% ethanol and processed for paraffin embedding. Sections at 6um thickness were cut and used for downstream staining procedures. Sections were stained with hematoxylin and eosin (H&E) to highlight the epithelium and potential post-infection related epithelial damage. For immunofluorescence staining, sections were allowed to dry in a 60° C for 1 hour. Sections were rehydrated in two washes of xylene for 5 min each followed by two washes in 100% EtOH (10 min each), 95% EtOH (10 min each), and H_2_O (5 min each). After rehydration, antigen retrieval was performed using sodium citrate buffer pH 6.0 or Tris-EDTA pH 9.0 in a pressure cooker for 30 minutes. Next, sections were blocked with 5% normal goat serum (005-000-121, Jackson ImmunoResearch) for 1 hour. Next, sections were incubated in the primary antibody overnight at 4° C at the indicated concentrations. The next day, sessions were stained with secondary antibodies for 2 hours at room temperature (RT) at the indicated concentrations. Finally, slides were stained with DAPI (5ug/ml; D9542, Sigma-Aldrich) for 5 minutes at RT and mounted in Prolong Gold Antifade Medium (P36930, Thermo Fisher Scientific). Primary antibodies were used at the following dilutions: rat-anti-E-cadherin (1:200; 13-1900, Thermo Fisher Scientific), rabbit-anti-Laminin (1:250; L9393, Sigma-Aldrich), mouse-anti-Ezrin (1:250; MA5-13862, Sigma-Aldrich), rabbit-anti-Ki67 (1:200; RM9106S0, Thermo Fischer Scientific), rabbit-anti-IFNγR1 (1:300; 10808-1-AP, Proteintech), rabbit-anti-IFNγR2 (1:300; 10266-1-AP, Proteintech). Secondary antibodies were used at the following dilutions: Alexa Fluor-conjugated secondary antibodies (1:500; A-21429, A-21245, A32728, Thermo Fisher Scientific), and rabbit biotinylated secondary antibodies (1:1000; BA-5000-1.5, Vector Laboratories). For immunostaining requiring signal amplification, TSA Cy3 Amplification kits were used (SAT704A001EA, PerkinElmer) and manufacturer-provided protocols were followed.

### Quantification of Lgr5 transcripts using RNAscope

RNA in situ hybridization for *Lgr5* expression was conducted on 7 μm paraffin sections using the RNAscope 2.5 High Definition (HD) – Red Assay (Advanced Cell Diagnostics, 322350). The manufacturer’s protocol was strictly followed, which included 15 minutes of target retrieval and 30 minutes of protease digestion, utilizing the RNAscope probe Mm-Lgr5 (Advanced Cell Diagnostics, 312171). Quantification of Lgr5 mRNA transcripts was carried out using the open-source platform Fiji ^75^, following the analysis guidelines provided by Advanced Cell Diagnostics. The area of individual probes was measured to determine the total probe count within probe clusters. Probe clusters containing at least 10 probes were quantified and normalized to the crypt area.

### Organoid culture media

Complete organoid 1 x ENR medium was prepared from 1 x Advanced DMEM/F12, 10 mM HEPES, 2 mM Glutamax, 0.11 mg/ml Penicillin-Streptomycin antibiotics, 1 mM N-Acetylcysteine, 50 ng/ml hEGF, 100ng/mlNoggin, 2%B-27 Supplement (ThermoFisher Scientific, 17504044), 1% N-2 supplement (Thermo Fisher Scientific, 17502048), and 250ng/ml of recombinant Rspo3 (rRspo3) (Genentech Inc) for culture of duodenum derived-organoids. Additionally, 10mM of Nicotinamide (7154; Stem Cell Technologies) and 250ng/ml of recombinant Wnt3a (1324-WN-010; R&D systems) were added in the above mentioned complete organoids 1 x ENR medium, creating a separate 1 x WENR medium which was used for culturing of colon-derived organoids. Both ENR and WENR were prepared fresh just prior to their use into the organoids during their establishment or passaging.

### Establishment of young and aged intestinal organoids from primary epithelial tissue

Organoid cultures were established from primary tissues as previously described ^76, 77^. Briefly, young and aged mice were sacrificed and dissected to harvest the intestine (duodenum and colon). Tissues were placed in 15ml of cold 1x PBS supplemented with 0.11mg/ml Penicillin-Streptomycin antibiotics, 2mM DTT, 1mM EDTA and 10uM Y-27632, and incubated on ice for 15 minutes. Next, intestines were moved to a tube with 20 ml cold PBS with 2mM DTT, 3mM EDTA, 10uM Y-27632 and incubated for an additional 60 minutes (duodenum) or 90 minutes (colon) followed by vigorous shaking for one minute to release crypts into the solution. Crypts were separated from villi material by filtering the solution using 70um cell strainers, followed by 2 washes with Advanced DMEM/F12 supplemented with 10mM HEPES, 0.11mg/ml Penicillin-Streptomycin antibiotics, and 1mM of N-Acetylcysteine. Finally, crypts were resuspended in Matrigel (356231, Corning), plated on 24-well culture plates and overlaid with 1x ENR (duodenum) or WENR (colon) medium to initiate organoid cultures (defining passage P0). Organoids were grown at 37° C in a 5% CO2 incubator, growth medium was changed at D5 and organoids were passaged at D7. For downstream experiments and analyses, organoids were cultured on 96-well culture plates.

### Establishment of fetal intestinal organoids from primary epithelial tissue

The protocol for dissecting mouse embryos, isolating fetal intestinal epithelial cells, and culturing intestinal spheroids was conducted as previously described^83^. Briefly, a pregnant C57BL/6 mouse (gestational day 15) was sacrificed, and the uterus was dissected out using dissection scissors. Embryos were isolated by making an incision along the uterus and separating Reichert’s membrane and visceral yolk sac, then gently transferred to a PBS dish. Under a stereomicroscope, an abdominal incision was made to open the peritoneal cavity, and the gastrointestinal organs, including the stomach, small intestine, cecum, colon, liver, and pancreas, were dissected out, isolating the small intestine. The small intestine was then transferred to a microcentrifuge tube and placed on ice. The intestine was chopped into pieces, incubated with 1 mM EDTA in PBS for 30 minutes on ice, and vortexed vigorously to separate epithelial cells. A small volume of the solution was checked under an inverted microscope to confirm epithelial cell detachment. The solution was then filtered through a 100 μm cell strainer, centrifuged at 200g for 5 minutes at 4°C, and the pellet was resuspended in 60 μL of cold Matrigel. The suspension was plated at 30 μL per well in a 24-well plate, incubated for 10 minutes at 37°C in a 5% CO2 incubator, and then 500 μL of culture medium was gently added to each well. Cells were cultured in a 37°C, 5% CO2 incubator for 5-7 days, with the medium replaced every other day and Y-27632 added for the first two days to prevent anoikis. For downstream experiments and analyses, organoids were cultured on 96-well culture plates.

### Mechanical passaging of young adult and aged intestinal organoids and treatments

Mechanical passaging of organoids was performed following established protocols ^78^. Briefly, on day 7 post-plating, Matrigel droplets containing organoids were disrupted by pipetting up and down 20–30 times using a P200 pipette and transferred to a 15 mL conical tube. The organoids were washed twice with 1x DMEM/F12 medium. The washed crypts were then replated at a 1:4 ratio into fresh Matrigel droplets and subsequently transferred into plastic 24-well plates for passaging or 96-well plates for downstream experiments (starting at P2).

For IFNγ treatments in colon derived-organoids, young and aged colonoids were passaged (P2) and at the same day (D0) were treated with 50ng/ml rIFNγ or PBS and were maintained in IFΝγ treatments for 60h. At this timepoint, all organoids were treated with CellEvent Caspase-3/7 Green ReadyProbes Reagent (R37111; ThermoFisher), according to the manufacturer’s protocol to identify apoptotic cells.

For IFNγ treatments in duodenum-derived young adult organoids, on D0 timepoint (same day as P2 passage), duodenum-derived young adult organoids were maintained in two separate culture conditions; ENR or WENR for 72h (D3) to induce either a fetal-like or an adult organoid phenotype. Next, organoids were treated with 20ng/ml rIFNγ or PBS for 48h. For blocking IFNγ-induced signaling activation in both fetal-like and adult organoids, they were treated with 20ng/ml rIFNγ and 10uM of Ruxolitinib for 48h. At this timepoint, all organoids were treated with CellEvent Caspase-3/7 Green ReadyProbes Reagent (R37111; ThermoFisher), according to the manufacturer’s protocol to identify apoptotic cells.

### Mechanical passaging of embryonic intestinal organoids and treatments

As previously described ^79^, to passage the embryonic organoids, the culture medium in the wells was first removed. Using a 1 mL pipettor, 1 mL of ice-cold 1 mM EDTA in PBS was added, and the solution was pipetted up and down vigorously to solubilize the Matrigel. The solution was then transferred into a 1.5 mL microcentrifuge tube and placed on ice. The spheroid suspension was pipetted up and down 20–30 times with P200 pipette tips to mechanically disrupt the spheroids into smaller fragments. The spheroid fragment suspension was centrifuged at 200g for 5 minutes at 4°C, and the supernatant was discarded. The pellet was resuspended in 90–150 μL of cold Matrigel solution, depending on the split ratio during passaging. A 30 μL aliquot of this solution was plated into the center of each well of a 24-well cell culture plate, which was then incubated for 10 minutes in a 37°C, 5% CO2 incubator. Subsequently, 500 μL of culture medium was gently added, and the cells were cultured in a 37°C, 5% CO2 incubator for 5–7 days. For IFNγ treatments, P2 fetal organoids were treated with 20ng/ml rIFNγ or PBS for 48h. At this timepoint, all organoids were treated with CellEvent Caspase-3/7 Green ReadyProbes Reagent (R37111; ThermoFisher), according to the manufacturer’s protocol to identify apoptotic cells.

### Image acquisition and analysis

Fluorescence and bright-field images of tissue sections were acquired using a Leica SP8 confocal microscope. Live imaged organoids were acquired using an incucyte S3 microscope. Stained organoids were acquired using a Leica SP8 confocal microscope. Images were processed with the open-source platform Fiji. Quantifications of immunofluorescence stainings were performed manually using Fiji. All tissue image analyses were performed in the distal 0.3 cm of the colons. Additionally, all quantifications were performed in a blind manner. For Ki67, a minimum of 15 well-oriented crypts were chosen randomly for quantification. Specifically, the number of Ki67+ cells as well as DAPI+ cells were quantified per crypt and based on these numbers, the % of Ki67+ cells per crypt was calculated. For Ezrin area quantification, a continuous line of the same thickness as Ezrin was drawn at the apical side of the epithelium encompassing a minimum of three 20x images (∼approximately 30-35 crypts) per sample. The area of the total line was measured and was considered the total apical area. Along that same total line, the area of that line that was also positively stained for Ezrin (defined as line that expressed signal>0 for Ezrin), was considered the Ezrin+ area. To get the % of Ezrin+ area, the Ezrin area was normalized by the total apical area. For Laminin quantification, the total mucosal area encompassing a minimum of three 20x images (∼approximately 30-35 crypts) per sample was quantified. From the same region, the Laminin+ area was also quantified. To get the % of Laminin+ area, the Laminin+ area was normalized by the total mucosal area. For IFNγR1 and R2 quantification, the total mucosal area encompassing a minimum of three 20x images (∼approximately 30-35 crypts) per sample was quantified. From the same region, the IFNγR1+ or IFNγR2+ area were also quantified. To get the % of IFNγR1+ or IFNγR2+ area, the IFNγR1+ or IFNγR2+ area was normalized by the total mucosal area. Regarding the organoid experiments, to quantify organoid survival, the number of initial seeded crypts at the start of an experiment was measured. Next, the number of alive organoids at the timepoint right before the end of the experiment was measured. From the above measurements, the % survival of organoids per condition was calculated. To quantify the MFI from CC3/7+ apoptotic organoids, the whole well was imaged (consisted of ∼ 30-50 organoids per condition). First, the background intensity was measured using Fiji (area of the well with no organoids). Next, the intensity of the whole well was measured and from this total intensity, the background intensity was subtracted and this was used as the MFI of a well. The same process was repeated for every single well of every different group. To calculate the MFI from Ly6a+, IFNγR1+, and IFNγR2+ organoids, first, the background intensity was measured using Fiji (area of the well with no organoids). Next, considering that these proteins are surface proteins, the protein intensity of the epithelium of each single organoid (a minimum of 10 well-oriented organoids were used per group) was quantified using Fiji. The background intensity was then subtracted from the organoid intensity and that final number was the organoid MFI. The average MFI from all organoids per well was used as the representative MFI per well. The same procedure was repeated for all different groups.

### Quantification and statistical analysis

Statistical analysis between groups was performed using GraphPad Prism 10 (GraphPad Software, La Jolla, CA). Value N represents the number of animals or independent organoid preparations. All graphs were first tested for normality distribution. Normally distributed data were analyzed using parametric Student’s t-test with Welch’s correction or one-way ANOVA with Tukey’s multiple comparisons test. The non-parametric Mann-Whitney U-test was used if the data did not fit a normal distribution. Significance was taken as P < 0.05 with a confidence interval of 95%. Data are presented as mean ± SD for parametric data or as median ± interquartile range for non-parametric data.

### Data and code availability

Bulk RNA-seq and scRNA-seq data have been deposited at GEO and are publicly available as of the date of publication. Accession numbers are listed in table S3. Microscopy data reported in this paper will be shared by the lead contacts upon request. Any additional information required to reanalyze the data reported in this paper is available from the lead contacts upon request.

## Author contributions

Conceptualization and design, A.K., H.T., J.D., R.P., D.G., H.J., D.C.-A; data generation, A.K., H.T., J.D., F.W., J.Y., Z.M., S.D., D.G., data analysis and interpretation, A.K., H.T., J.D., R.P., D.G., H.J., D.C.-A; writing-original draft, A.K., H.T., J.D., R.P., D.G., H.J., D.C.-A.

## Acknowledgments

We are grateful to Eliah Shamir, Susan Haller, Warrison Athanasio Coelho De Andrade, Dun Li for their thoughtful help, feedback, and suggestions on the manuscript. We are also grateful to Victor Nunez and Jessica Mills for necropsy support.

## Supplemental information

**Figure S1.**
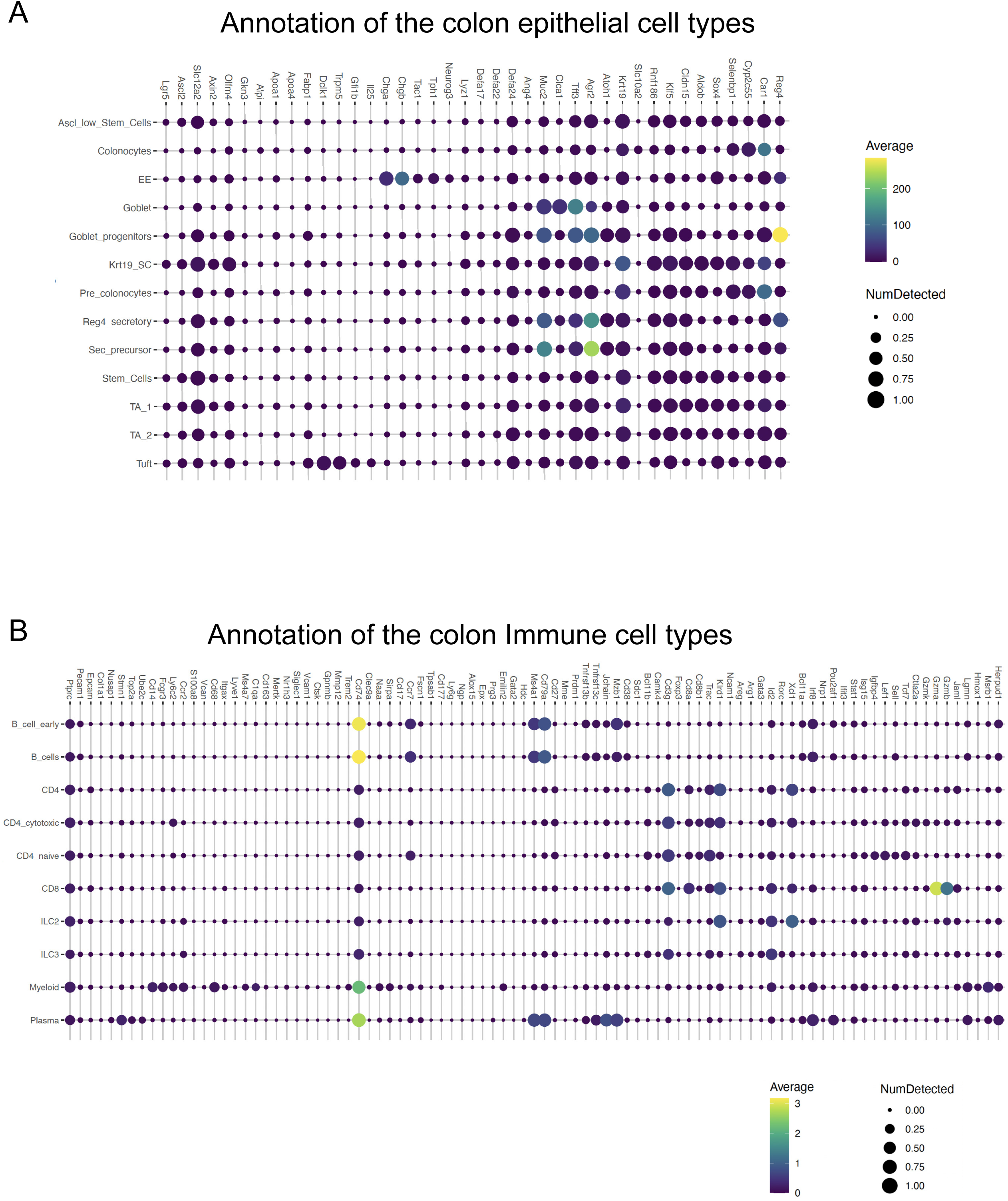
Markers used for annotation of the colon epithelial and immune cell types in young and aged mouse intestines. (A) Dotplot depicting the expression levels of the markers used for annotation of the colon epithelial cell types. (B) Dotplot depicting the expression levels of the markers used for annotation of the colon immune cell types..

**Figure S2.**
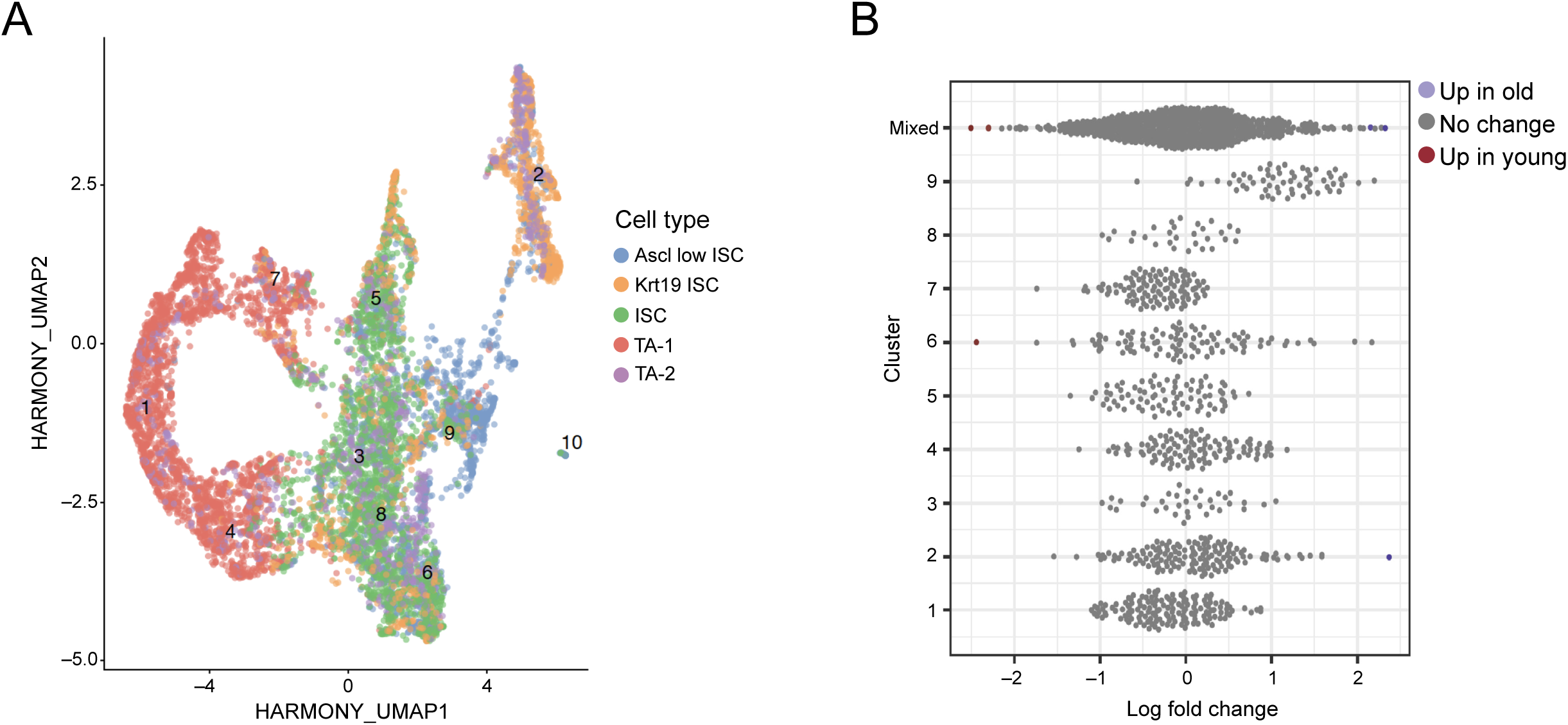
Composition analysis of young vs old stem cell and progenitor cells. (A) Sub-clustering of stem-cell like neighborhoods from the colon with per-sample level integration performed with Harmony. (B) Milo analysis of sub-clustered cells in (A). While very few neighborhoods reach nominal significance in a model containing a “study” effect, the Ascl2-low enriched cluster 9 shows a clear shift in estimated log FC in old colon relative to young. Colored dots represent significantly differentially abundant cell neighborhoods with a spatially adjusted FDR of < 0.1.

**Figure S3.**
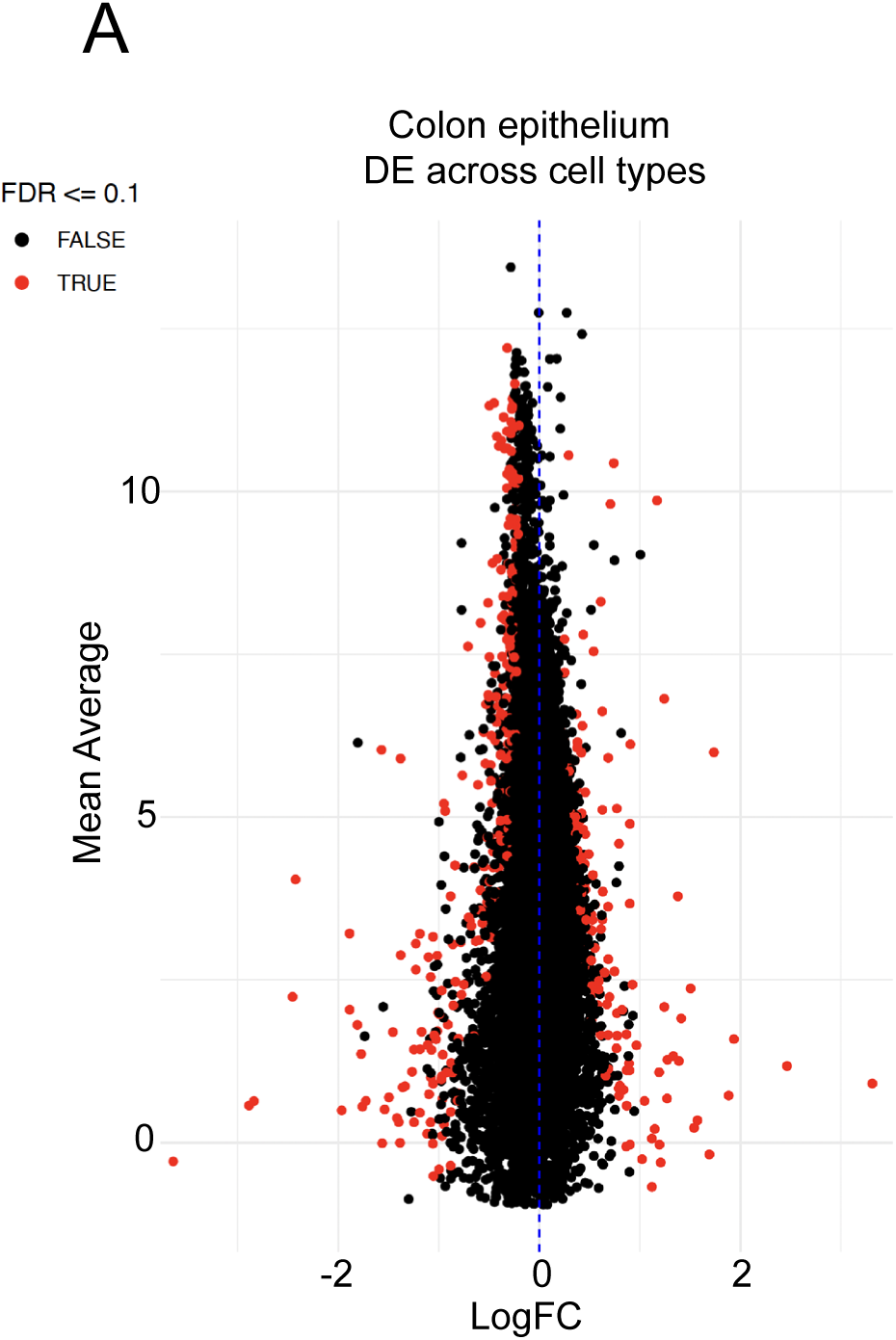
DE analysis of aged vs. young colon epithelium. (A) MA plot showing the average log2FC in gene expression between aged and young colon epithelium. Points shown in red pass an FDR threshold of <= 0.1.

**Figure S4.**
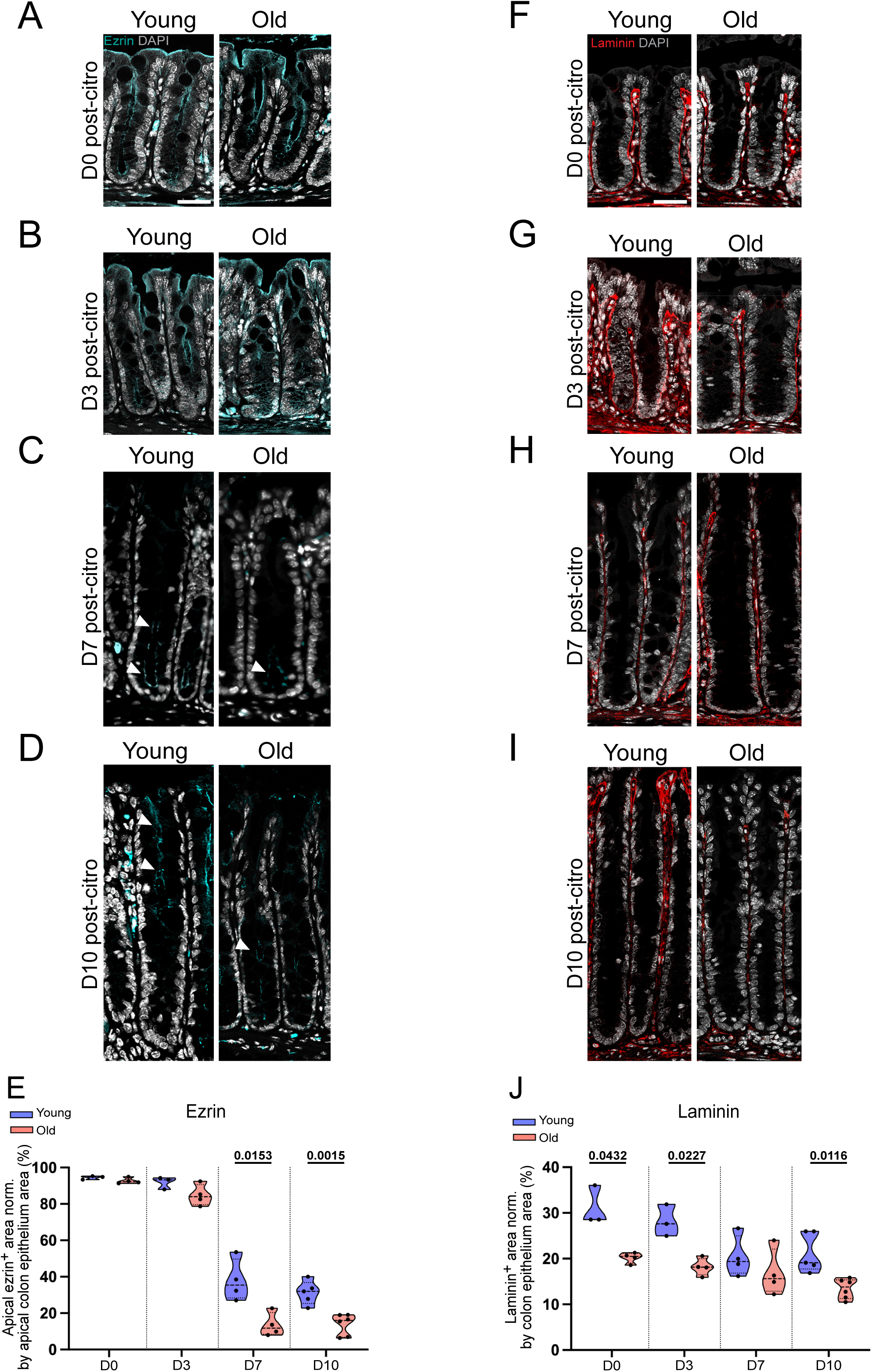
Aged colons exhibit higher epithelial damage during infection. (A-D) Representative Ezrin^+^ staining in young and aged colonic epithelium during homeostasis and infection (D0, 3, 7, and 10 post-*C. rodentium*). Scale bar: 40 μm. (E) Quantification of apical Ezrin^+^ area normalized by apical colon epithelium area from (A-D). Data are presented as mean and SD (3-6 biological samples per age group and infection timepoint were used for quantification). Statistical comparisons were performed using unpaired t-test with Welch’s correction (*P*≤0.05). (F-I) Representative Laminin^+^ staining in young and aged colonic epithelium during homeostasis and infection (D0, 3, 7, and 10 post-*C. rodentium*). Scale bar: 40 μm. (J) Quantification of Laminin^+^ area normalized by colon epithelium area from (F-I). Data are presented as mean and SD (3-6 biological samples per age group and infection timepoint were used for quantification). Statistical comparisons were performed using unpaired t-test with Welch’s correction (*P*≤0.05).

**Figure S5.**
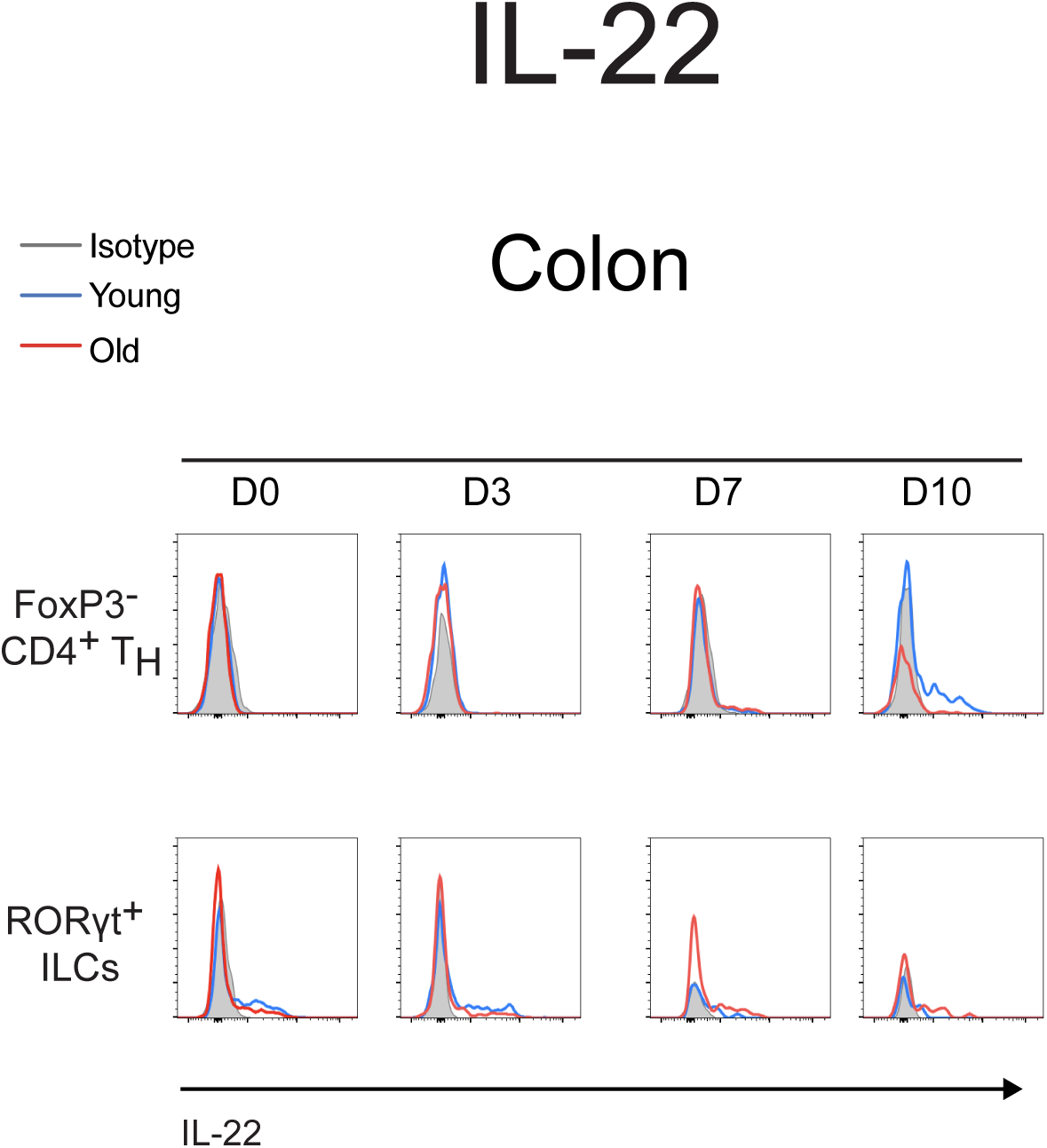
Decreased population of IL-22^+^ CD4^+^ T_H_ cells in the colon of old mice at day 10 post-*C. rodentium* infection. (A) Representative FACS plotting of young and aged IL22^+^ FoxP3^-^ CD4^+^ T_H_ and RORγt^+^ ILCs from colon during homeostasis and infection.

**Figure S6.**
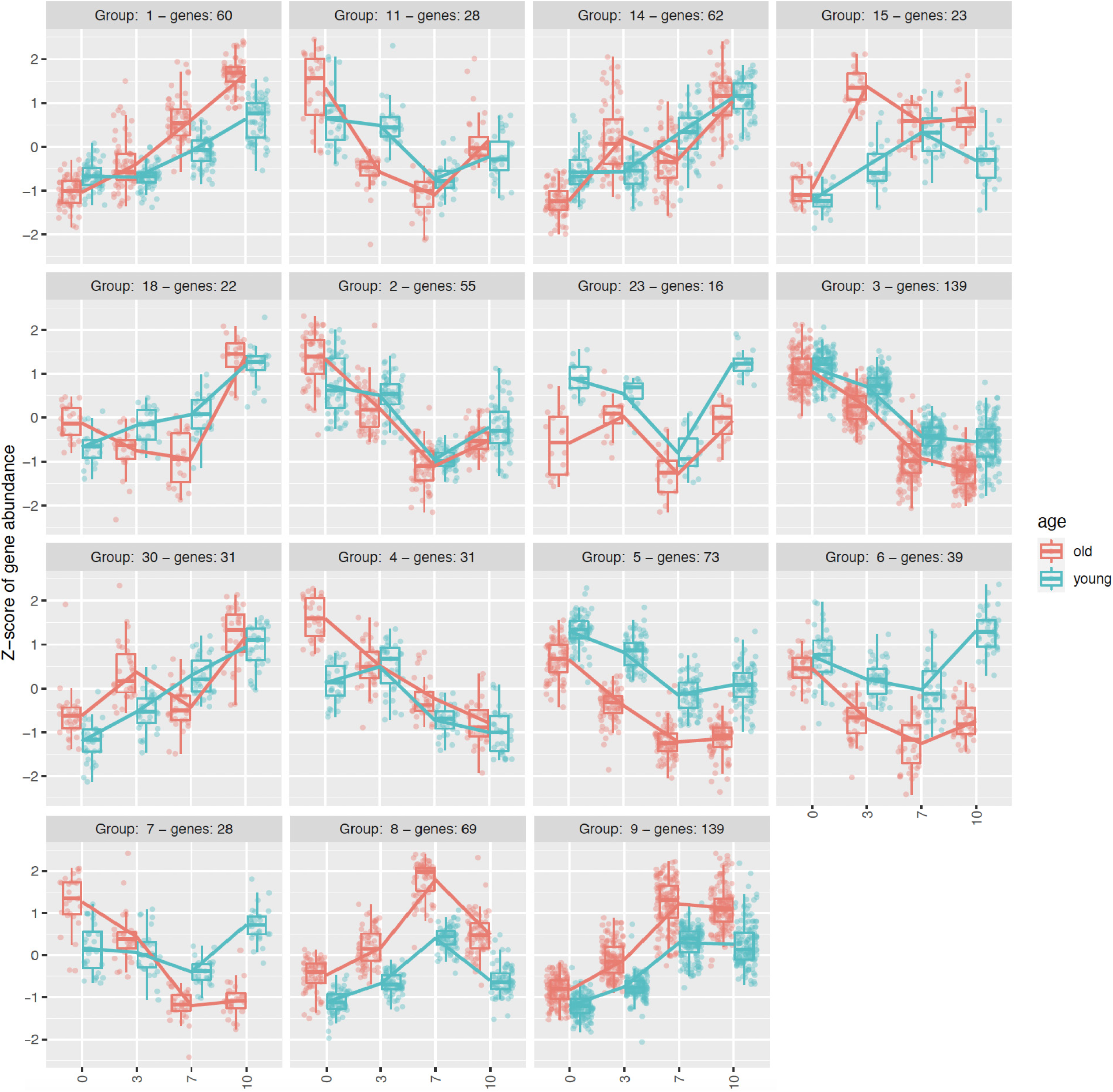
All clusters containing DE genes grouped based on their pattern of expression as a function of age.

**Figure S7.**
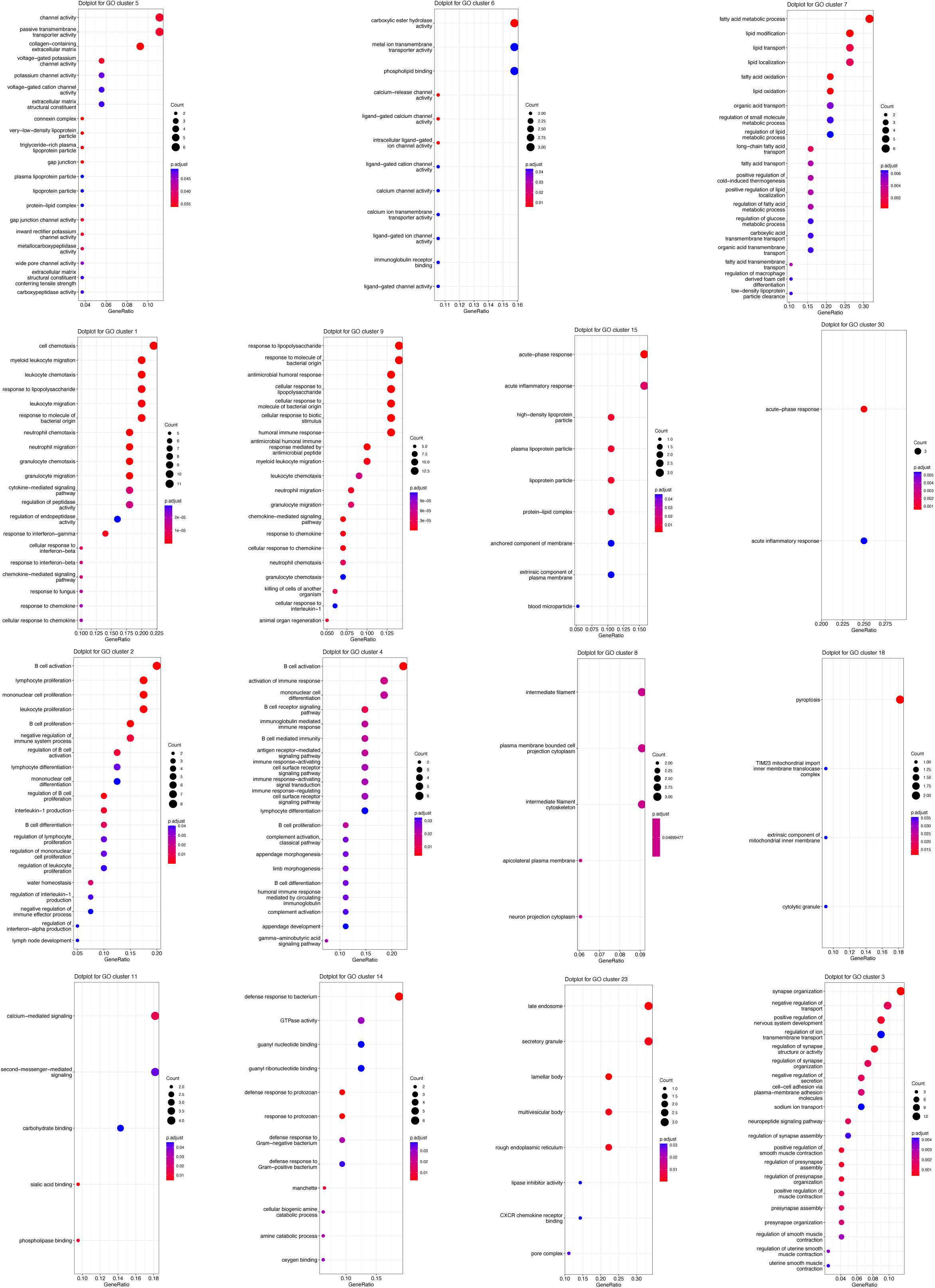
GO analysis of all clusters grouped based on their pattern of expression as a function of age.

**Figure S8.**
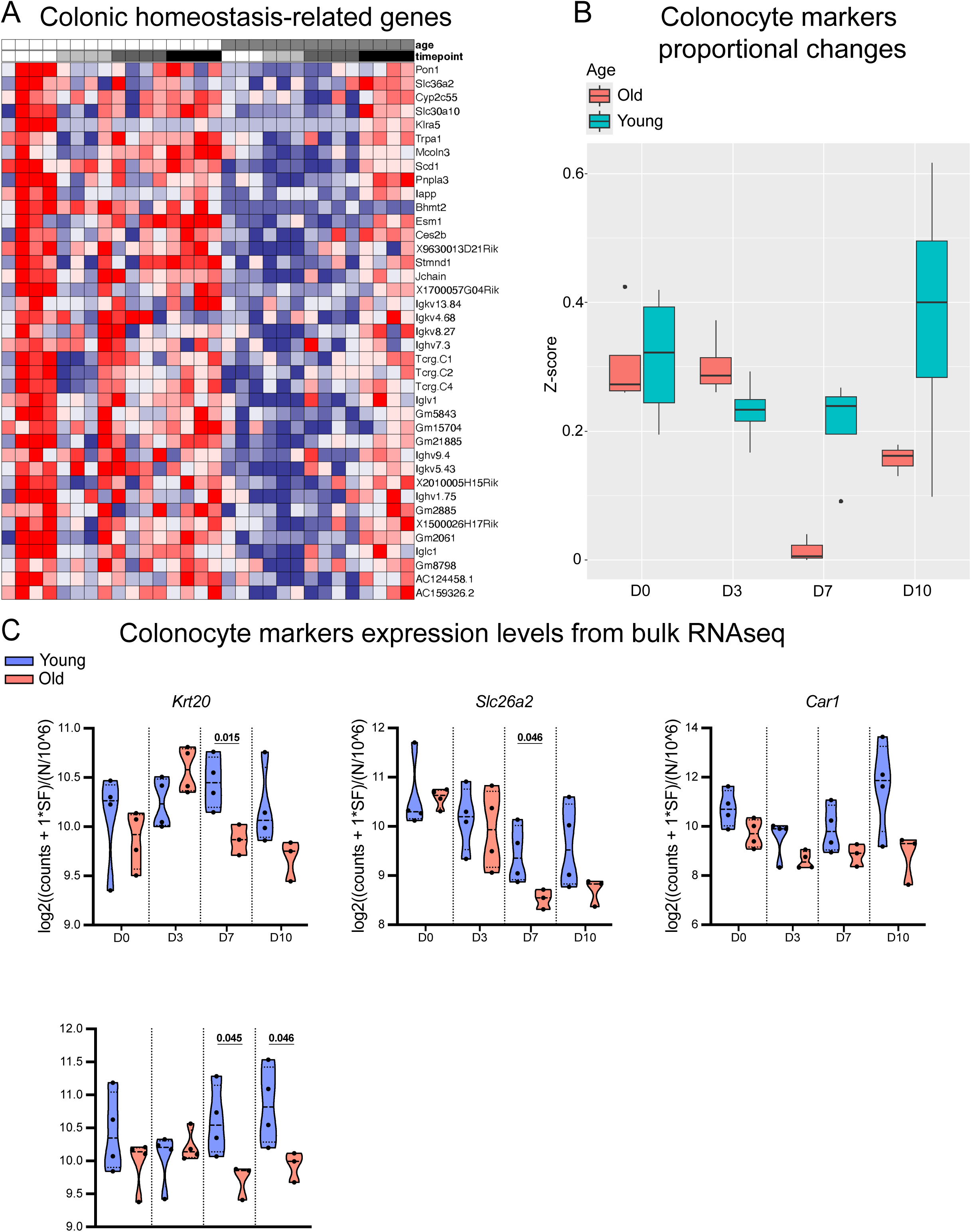
Decrease in colonic homeostasis genes and loss of colonocytes that are not replenished in aged infected colons. (A) Heatmap with all the DEGs from the colonic homeostasis-related cluster. (B) Estimated fraction of colonocyte cells based on the relative expression of colonocyte markers from a single cell reference (C) Expression levels of different colonocyte markers from bulk RNAseq in young and aged colons during homeostasis and infection. Data are presented as mean and SD (3-4 biological samples per age group and infection timepoint were used for quantification). Statistical comparisons were performed using unpaired t-test with Welch’s correction (*P*≤0.05).

**Figure S9.**
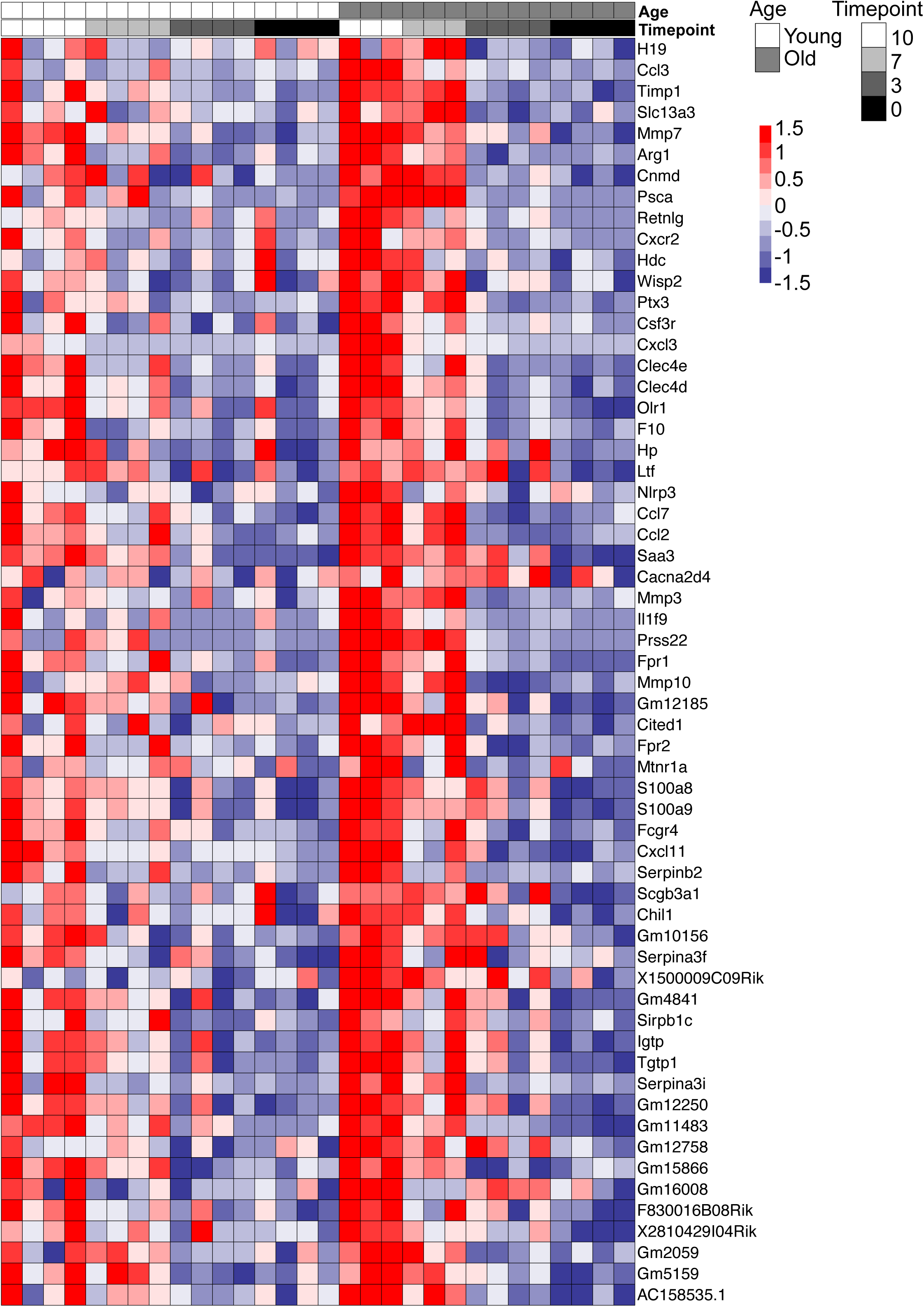
Expression of inflammatory-related genes increases by D7 in aged infected colons.

**Figure S10.**
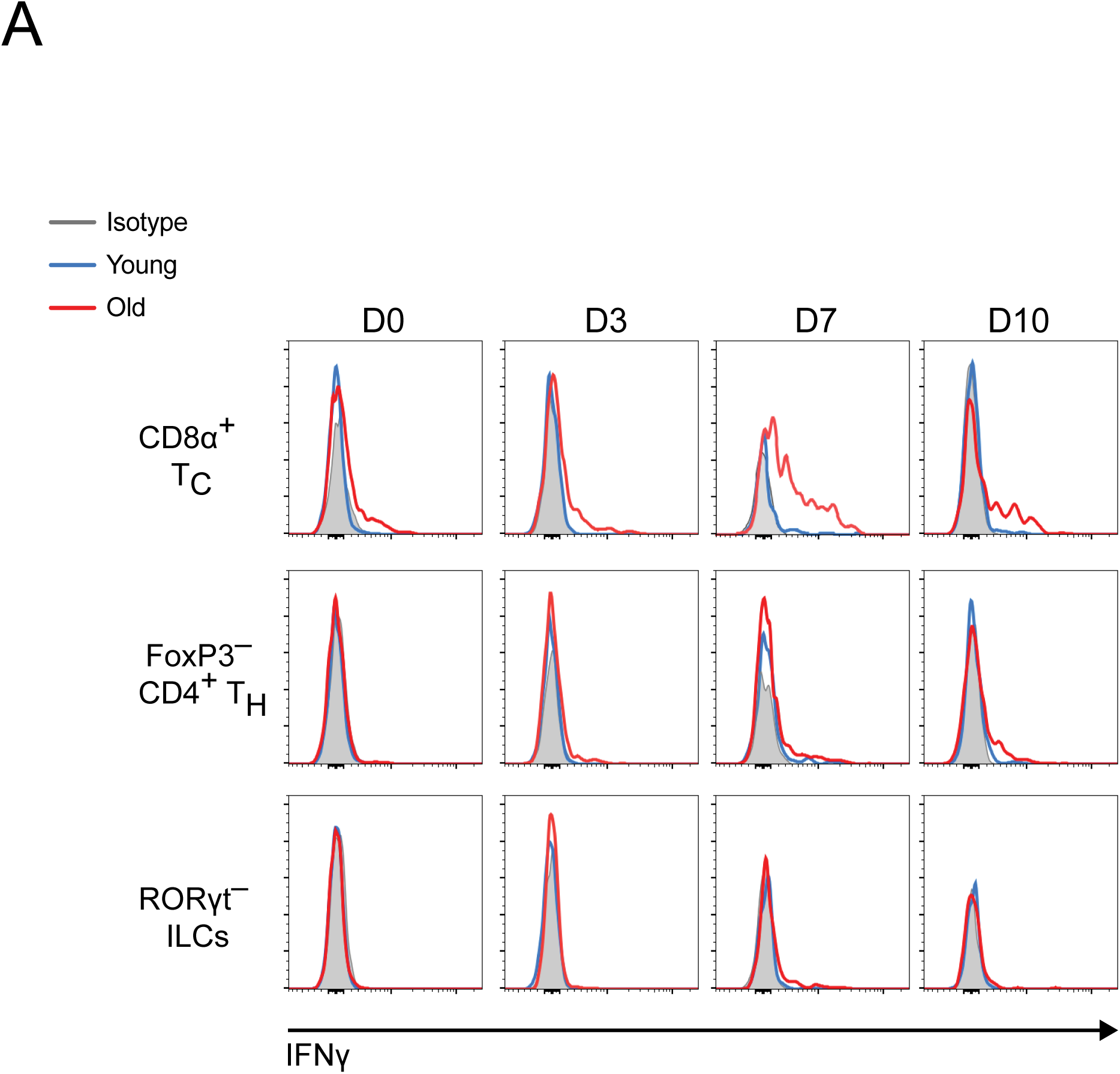
Frequency of IFNγ^+^ immune cells in the colon of *C. rodentium*-infected mice. (A) Representative FACS plotting of young and aged IFNγ^+^ CD8α^+^ Tc, FoxP3^-^ CD4^+^ T_H_, and RORγt^-^ ILCs from colon during homeostasis and infection.

**Figure S11.**
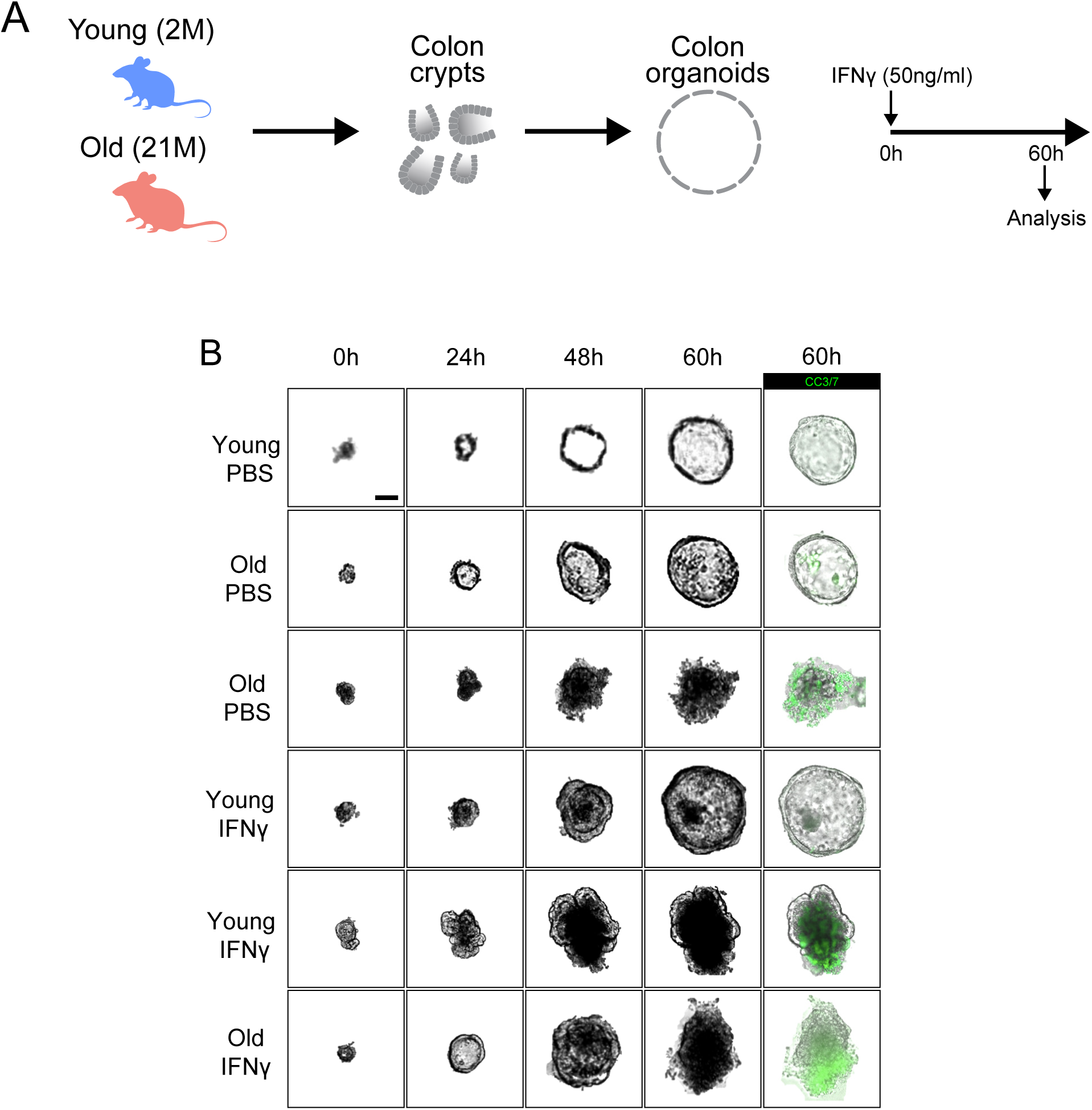
Aging enhances epithelial sensitivity to IFNγ and vulnerability to cell death. (A) Schematic of the experimental plan to assess the effect of aging and IFNγ in young and aged colon organoids. (B) Representative brightfield images of young and aged colon organoids. CC3/7 dye staining for assessment of apoptosis. Scale bar: 30 μm.

## References

1. Barnett, K., Mercer, S.W., Norbury, M., Watt, G., Wyke, S., and Guthrie, B. (2012). Epidemiology of multimorbidity and implications for health care, research, and medical education: a cross-sectional study. Lancet 380, 37–43. 10.1016/S0140-6736(12)60240-2.

2. Franceschi, C., Bonafe, M., Valensin, S., Olivieri, F., De Luca, M., Ottaviani, E., and De Benedictis, G. (2000). Inflamm-aging. An evolutionary perspective on immunosenescence. Ann N Y Acad Sci 908, 244–254. 10.1111/j.1749-6632.2000.tb06651.x.

3. Franceschi, C., Garagnani, P., Parini, P., Giuliani, C., and Santoro, A. (2018). Inflammaging: a new immune-metabolic viewpoint for age-related diseases. Nat Rev Endocrinol 14, 576–590. 10.1038/s41574-018-0059-4.

4. Fulop, T., Larbi, A., Pawelec, G., Khalil, A., Cohen, A.A., Hirokawa, K., Witkowski, J.M., and Franceschi, C. (2023). Immunology of Aging: the Birth of Inflammaging. Clin Rev Allergy Immunol 64, 109–122. 10.1007/s12016-021-08899-6.

5. Gavazzi, G., and Krause, K.H. (2002). Ageing and infection. Lancet Infect Dis 2, 659–666. 10.1016/s1473-3099(02)00437-1.

6. Pawelec, G., Goldeck, D., and Derhovanessian, E. (2014). Inflammation, ageing and chronic disease. Curr Opin Immunol 29, 23–28. 10.1016/j.coi.2014.03.007.

7. Nikolich-Zugich, J. (2018). The twilight of immunity: emerging concepts in aging of the immune system. Nat Immunol 19, 10–19. 10.1038/s41590-017-0006-x.

8. Pinti, M., Appay, V., Campisi, J., Frasca, D., Fulop, T., Sauce, D., Larbi, A., Weinberger, B., and Cossarizza, A. (2016). Aging of the immune system: Focus on inflammation and vaccination. Eur J Immunol 46, 2286–2301. 10.1002/eji.201546178.

9. Shaw, A.C., Goldstein, D.R., and Montgomery, R.R. (2013). Age-dependent dysregulation of innate immunity. Nat Rev Immunol 13, 875–887. 10.1038/nri3547.

10. Gehart, H., and Clevers, H. (2019). Tales from the crypt: new insights into intestinal stem cells. Nat Rev Gastroenterol Hepatol 16, 19–34. 10.1038/s41575-018-0081-y.

11. Castillo-Azofeifa, D., Fazio, E.N., Nattiv, R., Good, H.J., Wald, T., Pest, M.A., de Sauvage, F.J., Klein, O.D., and Asfaha, S. (2019). Atoh1(+) secretory progenitors possess renewal capacity independent of Lgr5(+) cells during colonic regeneration. EMBO J 38. 10.15252/embj.201899984.

12. Castillo-Azofeifa, D., Wald, T., Reyes, E.A., Gallagher, A., Schanin, J., Vlachos, S., Lamarche-Vane, N., Bomidi, C., Blutt, S., Estes, M.K., et al. (2023). A DLG1-ARHGAP31-CDC42 axis is essential for the intestinal stem cell response to fluctuating niche Wnt signaling. Cell Stem Cell 30, 188–206 e186. 10.1016/j.stem.2022.12.008.

13. Takashima, S., Martin, M.L., Jansen, S.A., Fu, Y., Bos, J., Chandra, D., O’Connor, M.H., Mertelsmann, A.M., Vinci, P., Kuttiyara, J., et al. (2019). T cell-derived interferon-gamma programs stem cell death in immune-mediated intestinal damage. Sci Immunol 4. 10.1126/sciimmunol.aay8556.

14. Biton, M., Haber, A.L., Rogel, N., Burgin, G., Beyaz, S., Schnell, A., Ashenberg, O., Su, C.W., Smillie, C., Shekhar, K., et al. (2018). T Helper Cell Cytokines Modulate Intestinal Stem Cell Renewal and Differentiation. Cell 175, 1307–1320 e1322. 10.1016/j.cell.2018.10.008.

15. Metidji, A., Omenetti, S., Crotta, S., Li, Y., Nye, E., Ross, E., Li, V., Maradana, M.R., Schiering, C., and Stockinger, B. (2018). The Environmental Sensor AHR Protects from Inflammatory Damage by Maintaining Intestinal Stem Cell Homeostasis and Barrier Integrity. Immunity 49, 353–362 e355. 10.1016/j.immuni.2018.07.010.

16. Becker, L., Nguyen, L., Gill, J., Kulkarni, S., Pasricha, P.J., and Habtezion, A. (2018). Age-dependent shift in macrophage polarisation causes inflammation-mediated degeneration of enteric nervous system. Gut 67, 827–836. 10.1136/gutjnl-2016-312940.

17. Nusse, Y.M., Savage, A.K., Marangoni, P., Rosendahl-Huber, A.K.M., Landman, T.A., de Sauvage, F.J., Locksley, R.M., and Klein, O.D. (2018). Parasitic helminths induce fetal-like reversion in the intestinal stem cell niche. Nature 559, 109–113. 10.1038/s41586-018-0257-1.

18. Yui, S., Azzolin, L., Maimets, M., Pedersen, M.T., Fordham, R.P., Hansen, S.L., Larsen, H.L., Guiu, J., Alves, M.R.P., Rundsten, C.F., et al. (2018). YAP/TAZ-Dependent Reprogramming of Colonic Epithelium Links ECM Remodeling to Tissue Regeneration. Cell Stem Cell 22, 35–49 e37. 10.1016/j.stem.2017.11.001.

19. Bala, P., Rennhack, J.P., Aitymbayev, D., Morris, C., Moyer, S.M., Duronio, G.N., Doan, P., Li, Z., Liang, X., Hornick, J.L., et al. (2023). Aberrant cell state plasticity mediated by developmental reprogramming precedes colorectal cancer initiation. Sci Adv 9, eadf0927. 10.1126/sciadv.adf0927.

20. Buschor, S., Cuenca, M., Uster, S.S., Scharen, O.P., Balmer, M.L., Terrazos, M.A., Schurch, C.M., and Hapfelmeier, S. (2017). Innate immunity restricts Citrobacter rodentium A/E pathogenesis initiation to an early window of opportunity. PLoS Pathog 13, e1006476. 10.1371/journal.ppat.1006476.

21. Silberger, D.J., Zindl, C.L., and Weaver, C.T. (2017). Citrobacter rodentium: a model enteropathogen for understanding the interplay of innate and adaptive components of type 3 immunity. Mucosal Immunol 10, 1108–1117. 10.1038/mi.2017.47.

22. Aparicio-Domingo, P., Romera-Hernandez, M., Karrich, J.J., Cornelissen, F., Papazian, N., Lindenbergh-Kortleve, D.J., Butler, J.A., Boon, L., Coles, M.C., Samsom, J.N., and Cupedo, T. (2015). Type 3 innate lymphoid cells maintain intestinal epithelial stem cells after tissue damage. J Exp Med 212, 1783–1791. 10.1084/jem.20150318.

23. Pickert, G., Neufert, C., Leppkes, M., Zheng, Y., Wittkopf, N., Warntjen, M., Lehr, H.A., Hirth, S., Weigmann, B., Wirtz, S., et al. (2009). STAT3 links IL-22 signaling in intestinal epithelial cells to mucosal wound healing. J Exp Med 206, 1465–1472. 10.1084/jem.20082683.

24. Lindemans, C.A., Calafiore, M., Mertelsmann, A.M., O’Connor, M.H., Dudakov, J.A., Jenq, R.R., Velardi, E., Young, L.F., Smith, O.M., Lawrence, G., et al. (2015). Interleukin-22 promotes intestinal-stem-cell-mediated epithelial regeneration. Nature 528, 560–564. 10.1038/nature16460.

25. Zenewicz, L.A., Yancopoulos, G.D., Valenzuela, D.M., Murphy, A.J., Stevens, S., and Flavell, R.A. (2008). Innate and adaptive interleukin-22 protects mice from inflammatory bowel disease. Immunity 29, 947–957. 10.1016/j.immuni.2008.11.003.

26. Zha, J.M., Li, H.S., Lin, Q., Kuo, W.T., Jiang, Z.H., Tsai, P.Y., Ding, N., Wu, J., Xu, S.F., Wang, Y.T., et al. (2019). Interleukin 22 Expands Transit-Amplifying Cells While Depleting Lgr5(+) Stem Cells via Inhibition of Wnt and Notch Signaling. Cell Mol Gastroenterol Hepatol 7, 255–274. 10.1016/j.jcmgh.2018.09.006.

27. Mahapatro, M., Erkert, L., and Becker, C. (2021). Cytokine-Mediated Crosstalk between Immune Cells and Epithelial Cells in the Gut. Cells 10. 10.3390/cells10010111.

28. Omrani, O., Krepelova, A., Rasa, S.M.M., Sirvinskas, D., Lu, J., Annunziata, F., Garside, G., Bajwa, S., Reinhardt, S., Adam, L., et al. (2023). Author Correction: IFNgamma-Stat1 axis drives aging-associated loss of intestinal tissue homeostasis and regeneration. Nat Commun 14, 6302. 10.1038/s41467-023-42168-8.

29. Funk, M.C., Gleixner, J.G., Heigwer, F., Vonficht, D., Valentini, E., Aydin, Z., Tonin, E., Del Prete, S., Mahara, S., Throm, Y., et al. (2023). Aged intestinal stem cells propagate cell-intrinsic sources of inflammaging in mice. Dev Cell 58, 2914–2929 e2917. 10.1016/j.devcel.2023.11.013.

30. Mihaylova, M.M., Cheng, C.W., Cao, A.Q., Tripathi, S., Mana, M.D., Bauer-Rowe, K.E., Abu-Remaileh, M., Clavain, L., Erdemir, A., Lewis, C.A., et al. (2018). Fasting Activates Fatty Acid Oxidation to Enhance Intestinal Stem Cell Function during Homeostasis and Aging. Cell Stem Cell 22, 769–778 e764. 10.1016/j.stem.2018.04.001.

31. Pentinmikko, N., Iqbal, S., Mana, M., Andersson, S., Cognetta, A.B., 3rd, Suciu, R.M., Roper, J., Luopajarvi, K., Markelin, E., Gopalakrishnan, S., et al. (2019). Notum produced by Paneth cells attenuates regeneration of aged intestinal epithelium. Nature 571, 398–402. 10.1038/s41586-019-1383-0.

32. Nalapareddy, K., Nattamai, K.J., Kumar, R.S., Karns, R., Wikenheiser-Brokamp, K.A., Sampson, L.L., Mahe, M.M., Sundaram, N., Yacyshyn, M.B., Yacyshyn, B., et al. (2017). Canonical Wnt Signaling Ameliorates Aging of Intestinal Stem Cells. Cell Rep 18, 2608–2621. 10.1016/j.celrep.2017.02.056.

33. Gebert, N., Cheng, C.W., Kirkpatrick, J.M., Di Fraia, D., Yun, J., Schadel, P., Pace, S., Garside, G.B., Werz, O., Rudolph, K.L., et al. (2020). Region-Specific Proteome Changes of the Intestinal Epithelium during Aging and Dietary Restriction. Cell Rep 31, 107565. 10.1016/j.celrep.2020.107565.

34. Omrani, O., Krepelova, A., Rasa, S.M.M., Sirvinskas, D., Lu, J., Annunziata, F., Garside, G., Bajwa, S., Reinhardt, S., Adam, L., et al. (2023). IFNgamma-Stat1 axis drives aging-associated loss of intestinal tissue homeostasis and regeneration. Nat Commun 14, 6109. 10.1038/s41467-023-41683-y.

35. Liu, A., Lv, H., Wang, H., Yang, H., Li, Y., and Qian, J. (2020). Aging Increases the Severity of Colitis and the Related Changes to the Gut Barrier and Gut Microbiota in Humans and Mice. J Gerontol A Biol Sci Med Sci 75, 1284–1292. 10.1093/gerona/glz263.

36. Liu, Y., Ji, Y., Jiang, R., Fang, C., Shi, G., Cheng, L., Zuo, Y., Ye, Y., Su, X., Li, J., et al. (2023). Reduced smooth muscle-fibroblasts transformation potentially decreases intestinal wound healing and colitis-associated cancer in ageing mice. Signal Transduct Target Ther 8, 294. 10.1038/s41392-023-01554-w.

37. Jo, M.K., Moon, C.M., Jeon, H.J., Han, Y., Lee, E.S., Kwon, J.H., Yang, K.M., Ahn, Y.H., Kim, S.E., Jung, S.A., and Kim, T.I. (2023). Effect of aging on the formation and growth of colonic epithelial organoids by changes in cell cycle arrest through TGF-beta-Smad3 signaling. Inflamm Regen 43, 35. 10.1186/s41232-023-00282-6.

38. Kobayashi, T., Siegmund, B., Le Berre, C., Wei, S.C., Ferrante, M., Shen, B., Bernstein, C.N., Danese, S., Peyrin-Biroulet, L., and Hibi, T. (2020). Ulcerative colitis. Nat Rev Dis Primers 6, 74. 10.1038/s41572-020-0205-x.

39. Roda, G., Chien Ng, S., Kotze, P.G., Argollo, M., Panaccione, R., Spinelli, A., Kaser, A., Peyrin-Biroulet, L., and Danese, S. (2020). Crohn’s disease. Nat Rev Dis Primers 6, 22. 10.1038/s41572-020-0156-2.

40. Tursi, A., Scarpignato, C., Strate, L.L., Lanas, A., Kruis, W., Lahat, A., and Danese, S. (2020). Colonic diverticular disease. Nat Rev Dis Primers 6, 20. 10.1038/s41572-020-0153-5.

41. Barsouk, A., Rawla, P., Barsouk, A., and Thandra, K.C. (2019). Epidemiology of Cancers of the Small Intestine: Trends, Risk Factors, and Prevention. Med Sci (Basel) 7. 10.3390/medsci7030046.

42. Sartor, R.B. (2008). Microbial influences in inflammatory bowel diseases. Gastroenterology 134, 577–594. 10.1053/j.gastro.2007.11.059.

43. Guarner, F., and Malagelada, J.R. (2003). Gut flora in health and disease. Lancet 361, 512–519. 10.1016/S0140-6736(03)12489-0.

44. Sirvinskas, D., Omrani, O., Lu, J., Rasa, M., Krepelova, A., Adam, L., Kaeppel, S., Sommer, F., and Neri, F. (2022). Single-cell atlas of the aging mouse colon. iScience 25, 104202. 10.1016/j.isci.2022.104202.

45. Dann, E., Henderson, N.C., Teichmann, S.A., Morgan, M.D., and Marioni, J.C. (2022). Differential abundance testing on single-cell data using k-nearest neighbor graphs. Nat Biotechnol 40, 245–253. 10.1038/s41587-021-01033-z.

46. Subramanian, A., Tamayo, P., Mootha, V.K., Mukherjee, S., Ebert, B.L., Gillette, M.A., Paulovich, A., Pomeroy, S.L., Golub, T.R., Lander, E.S., and Mesirov, J.P. (2005). Gene set enrichment analysis: a knowledge-based approach for interpreting genome-wide expression profiles. Proc Natl Acad Sci U S A 102, 15545–15550. 10.1073/pnas.0506580102.

47. Castanza, A.S., Recla, J.M., Eby, D., Thorvaldsdottir, H., Bult, C.J., and Mesirov, J.P. (2023). Extending support for mouse data in the Molecular Signatures Database (MSigDB). Nat Methods 20, 1619–1620. 10.1038/s41592-023-02014-7.

48. Geremia, A., and Arancibia-Carcamo, C.V. (2017). Innate Lymphoid Cells in Intestinal Inflammation. Front Immunol 8, 1296. 10.3389/fimmu.2017.01296.

49. Amadou Amani, S., and Lang, M.L. (2020). Bacteria That Cause Enteric Diseases Stimulate Distinct Humoral Immune Responses. Front Immunol 11, 565648. 10.3389/fimmu.2020.565648.

50. Cox, C.B., Storm, E.E., Kapoor, V.N., Chavarria-Smith, J., Lin, D.L., Wang, L., Li, Y., Kljavin, N., Ota, N., Bainbridge, T.W., et al. (2021). IL-1R1-dependent signaling coordinates epithelial regeneration in response to intestinal damage. Sci Immunol 6. 10.1126/sciimmunol.abe8856.

51. Li, A.C., and Thompson, R.P. (2003). Basement membrane components. J Clin Pathol 56, 885–887. 10.1136/jcp.56.12.885.

52. Casaletto, J.B., Saotome, I., Curto, M., and McClatchey, A.I. (2011). Ezrin-mediated apical integrity is required for intestinal homeostasis. Proc Natl Acad Sci U S A 108, 11924–11929. 10.1073/pnas.1103418108.

53. Hopkins, E.G.D., Roumeliotis, T.I., Mullineaux-Sanders, C., Choudhary, J.S., and Frankel, G. (2019). Intestinal Epithelial Cells and the Microbiome Undergo Swift Reprogramming at the Inception of Colonic Citrobacter rodentium Infection. mBio 10. 10.1128/mBio.00062-19.

54. Zindl, C.L., Garrett Wilson, C., Chadha, A.S., Cai, B., Harbour, S.N., Nagaoka-Kamata, Y., Hatton, R.D., Gao, M., Figge, D.A., and Weaver, C.T. (2023). 10.1101/2023.05.03.539269.

55. Wang, L., Wang, E., Wang, Y., Mines, R., Xiang, K., Sun, Z., Zhou, G., Chen, K.Y., Rakhilin, N., Chao, S., et al. (2018). miR-34a is a microRNA safeguard for Citrobacter-induced inflammatory colon oncogenesis. Elife 7. 10.7554/eLife.39479.

56. Mullineaux-Sanders, C., Kozik, Z., Sanchez-Garrido, J., Hopkins, E.G.D., Choudhary, J.S., and Frankel, G. (2021). Citrobacter rodentium Infection Induces Persistent Molecular Changes and Interferon Gamma-Dependent Major Histocompatibility Complex Class II Expression in the Colonic Epithelium. mBio 13, e0323321. 10.1128/mbio.03233-21.

57. Zwarycz, B., Gracz, A.D., Rivera, K.R., Williamson, I.A., Samsa, L.A., Starmer, J., Daniele, M.A., Salter-Cid, L., Zhao, Q., and Magness, S.T. (2019). IL22 Inhibits Epithelial Stem Cell Expansion in an Ileal Organoid Model. Cell Mol Gastroenterol Hepatol 7, 1–17. 10.1016/j.jcmgh.2018.06.008.

58. Pantano, L. (2023). DEGreport: Report of DEG analysis. https://bioconductor.org/packages/DEGreport/.

59. Lu, X., Luo, C., Wu, J., Deng, Y., Mu, X., Zhang, T., Yang, X., Liu, Q., Li, Z., Tang, S., et al. (2023). Ion channels and transporters regulate nutrient absorption in health and disease. J Cell Mol Med 27, 2631–2642. 10.1111/jcmm.17853.

60. van der Lugt, B., Rusli, F., Lute, C., Lamprakis, A., Salazar, E., Boekschoten, M.V., Hooiveld, G.J., Muller, M., Vervoort, J., Kersten, S., et al. (2018). Integrative analysis of gut microbiota composition, host colonic gene expression and intraluminal metabolites in aging C57BL/6J mice. Aging (Albany NY) 10, 930–950. 10.18632/aging.101439.

61. Eriguchi, Y., Nakamura, K., Yokoi, Y., Sugimoto, R., Takahashi, S., Hashimoto, D., Teshima, T., Ayabe, T., Selsted, M.E., and Ouellette, A.J. (2018). Essential role of IFN-gamma in T cell-associated intestinal inflammation. JCI Insight 3. 10.1172/jci.insight.121886.

62. Merenda, A., Fenderico, N., and Maurice, M.M. (2020). Wnt Signaling in 3D: Recent Advances in the Applications of Intestinal Organoids. Trends Cell Biol 30, 60–73. 10.1016/j.tcb.2019.10.003.

63. Viragova, S., Li, D., and Klein, O.D. (2024). Activation of fetal-like molecular programs during regeneration in the intestine and beyond. Cell Stem Cell 31, 949–960. 10.1016/j.stem.2024.05.009.

64. Iqbal, S., Andersson, S., Nestaite, E., Pentinmikko, N., Kumar, A., Borshagovski, D., Webb, A., Saarinen, T., Juuti, A., Ori, A., et al. (2021). 10.1101/2021.06.24.449590.

65. Nava, P., Koch, S., Laukoetter, M.G., Lee, W.Y., Kolegraff, K., Capaldo, C.T., Beeman, N., Addis, C., Gerner-Smidt, K., Neumaier, I., et al. (2010). Interferon-gamma regulates intestinal epithelial homeostasis through converging beta-catenin signaling pathways. Immunity 32, 392–402. 10.1016/j.immuni.2010.03.001.

66. Raetz, M., Hwang, S.H., Wilhelm, C.L., Kirkland, D., Benson, A., Sturge, C.R., Mirpuri, J., Vaishnava, S., Hou, B., Defranco, A.L., et al. (2013). Parasite-induced TH1 cells and intestinal dysbiosis cooperate in IFN-gamma-dependent elimination of Paneth cells. Nat Immunol 14, 136–142. 10.1038/ni.2508.

67. Farin, H.F., Karthaus, W.R., Kujala, P., Rakhshandehroo, M., Schwank, G., Vries, R.G., Kalkhoven, E., Nieuwenhuis, E.E., and Clevers, H. (2014). Paneth cell extrusion and release of antimicrobial products is directly controlled by immune cell-derived IFN-gamma. J Exp Med 211, 1393–1405. 10.1084/jem.20130753.

68. Oost, K.C., Kahnwald, M., Barbiero, S., de Medeiros, G., Suppinger, S., Kalck, V., Meylan, L.C., Smallwood, S.A., Stadler, M.B., and Liberali, P. (2023). 10.1101/2023.12.18.572103.

69. Serra, D., Mayr, U., Boni, A., Lukonin, I., Rempfler, M., Challet Meylan, L., Stadler, M.B., Strnad, P., Papasaikas, P., Vischi, D., et al. (2019). Self-organization and symmetry breaking in intestinal organoid development. Nature 569, 66–72. 10.1038/s41586-019-1146-y.

70. Haghverdi, L., Lun, A.T.L., Morgan, M.D., and Marioni, J.C. (2018). Batch effects in single-cell RNA-sequencing data are corrected by matching mutual nearest neighbors. Nat Biotechnol 36, 421–427. 10.1038/nbt.4091.

71. Hao, Y., Hao, S., Andersen-Nissen, E., Mauck, W.M., 3rd, Zheng, S., Butler, A., Lee, M.J., Wilk, A.J., Darby, C., Zager, M., et al. (2021). Integrated analysis of multimodal single-cell data. Cell 184, 3573–3587 e3529. 10.1016/j.cell.2021.04.048.

72. Stuart, T., Butler, A., Hoffman, P., Hafemeister, C., Papalexi, E., Mauck, W.M., 3rd, Hao, Y., Stoeckius, M., Smibert, P., and Satija, R. (2019). Comprehensive Integration of Single-Cell Data. Cell 177, 1888–1902 e1821. 10.1016/j.cell.2019.05.031.

73. Law, C.W., Chen, Y., Shi, W., and Smyth, G.K. (2014). voom: Precision weights unlock linear model analysis tools for RNA-seq read counts. Genome Biol 15, R29. 10.1186/gb-2014-15-2-r29.

74. Ritchie, M.E., Phipson, B., Wu, D., Hu, Y., Law, C.W., Shi, W., and Smyth, G.K. (2015). limma powers differential expression analyses for RNA-sequencing and microarray studies. Nucleic Acids Res 43, e47. 10.1093/nar/gkv007.

75. Schindelin, J., Arganda-Carreras, I., Frise, E., Kaynig, V., Longair, M., Pietzsch, T., Preibisch, S., Rueden, C., Saalfeld, S., Schmid, B., et al. (2012). Fiji: an open-source platform for biological-image analysis. Nat Methods 9, 676–682. 10.1038/nmeth.2019.

76. Sanman, L.E., Chen, I.W., Bieber, J.M., Steri, V., Trentesaux, C., Hann, B., Klein, O.D., Wu, L.F., and Altschuler, S.J. (2021). Transit-Amplifying Cells Coordinate Changes in Intestinal Epithelial Cell-Type Composition. Dev Cell 56, 356–365 e359. 10.1016/j.devcel.2020.12.020.

77. Sanman, L.E., Chen, I.W., Bieber, J.M., Thorne, C.A., Wu, L.F., and Altschuler, S.J. (2020). Generation and Quantitative Imaging of Enteroid Monolayers. Methods Mol Biol 2171, 99–113. 10.1007/978-1-0716-0747-3_6.

78. Sato, T., Vries, R.G., Snippert, H.J., van de Wetering, M., Barker, N., Stange, D.E., van Es, J.H., Abo, A., Kujala, P., Peters, P.J., and Clevers, H. (2009). Single Lgr5 stem cells build crypt-villus structures in vitro without a mesenchymal niche. Nature 459, 262–265. 10.1038/nature07935.

79. Imajo, M., Hirota, A., and Tanaka, S. (2023). Generation of Fetal Intestinal Organoids and Their Maturation into Adult Intestinal Cells. Methods Mol Biol 2650, 133–140. 10.1007/978-1-0716-3076-1_11.

